# The MRE11-RAD50-NBS1 complex both starts and extends DNA end resection in mouse meiosis

**DOI:** 10.1101/2024.08.17.608390

**Authors:** Soonjoung Kim, Shintaro Yamada, Tao Li, Claudia Canasto-Chibuque, Jun Hyun Kim, Marina Marcet-Ortega, Jiaqi Xu, Diana Y. Eng, Laura Feeney, John H. J. Petrini, Scott Keeney

## Abstract

Nucleolytic resection of DNA ends is critical for homologous recombination, but its mechanism is not fully understood, particularly in mammalian meiosis. Here we examine roles of the conserved MRN complex (MRE11, RAD50, and NBS1) through genome-wide analysis of meiotic resection in mice with various MRN mutations, including several that cause chromosomal instability in humans. Meiotic DSBs form at elevated levels but remain unresected if *Mre11* is conditionally deleted, thus MRN is required for both resection initiation and regulation of DSB numbers. Resection lengths are reduced to varying degrees in MRN hypomorphs or if MRE11 nuclease activity is attenuated in a conditional nuclease-dead *Mre11* model. These findings unexpectedly establish that MRN is needed for longer-range extension of resection, not just resection initiation. Finally, resection defects are additively worsened by combining MRN and *Exo1* mutations, and mice that are unable to initiate resection or have greatly curtailed resection lengths experience catastrophic spermatogenic failure. Our results elucidate multiple functions of MRN in meiotic recombination, uncover unanticipated relationships between short- and long-range resection, and establish the importance of resection for mammalian meiosis.

## INTRODUCTION

Homologous recombination during meiosis initiates with DNA double-strand breaks (DSBs) made by SPO11 protein, which remains covalently bound to DNA 5’ ends after strand breakage^1,2^. These DSBs must then be nucleolytically processed (resected) to generate the single-stranded DNA (ssDNA) needed for homology search and strand exchange (**Figure 1A**)^3^. Despite its central role in recombination, meiotic DSB resection remains poorly understood, particularly in mammals.

**Figure 1.**
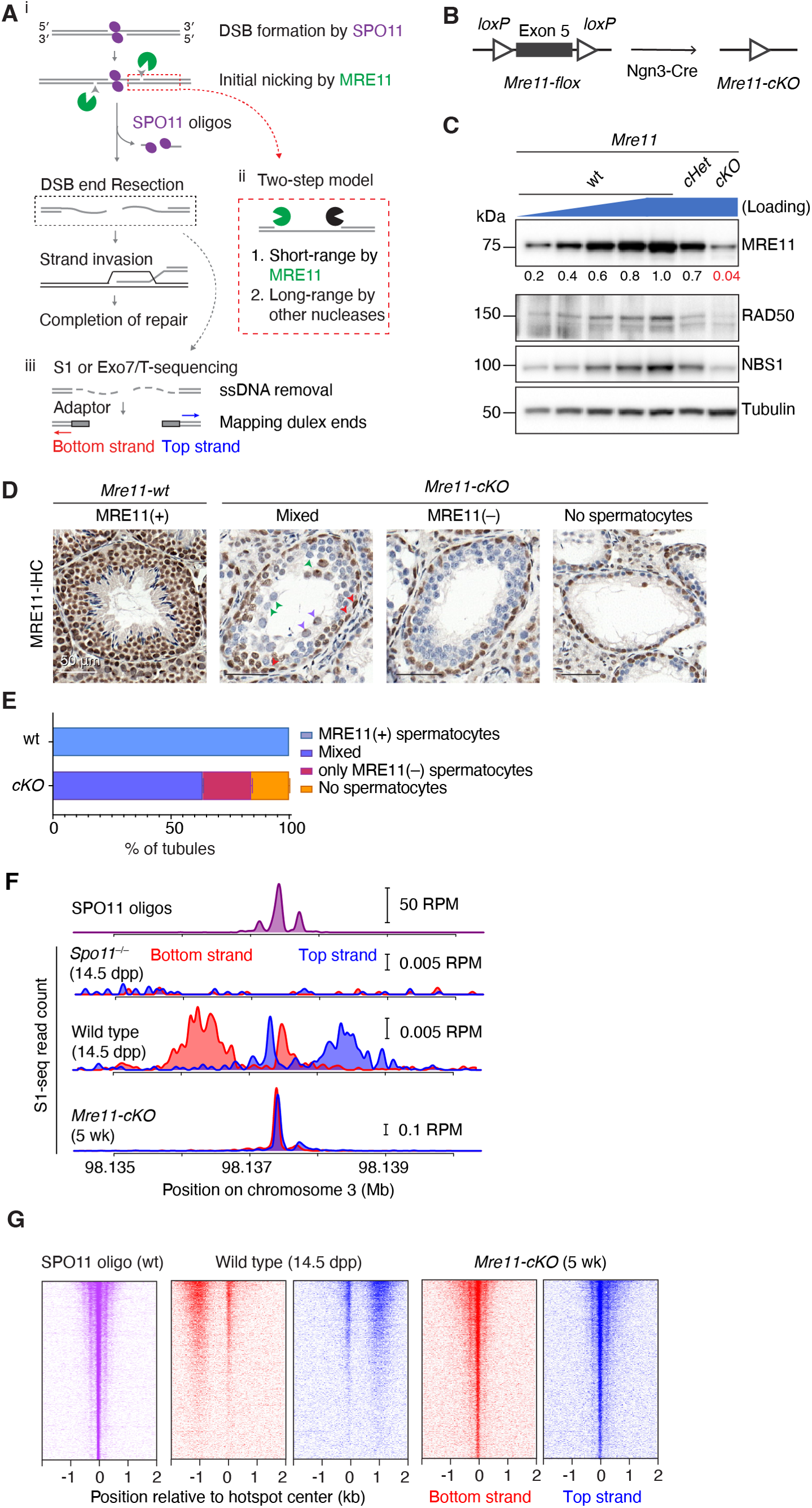
MRE11 is required for resection initiation. (A) Early meiotic recombination steps and sequencing strategy. i, SPO11 (magenta ellipses) cuts DNA via a covalent protein–DNA intermediate. SPO11-bound strands are nicked (arrowheads) to give an entry point(s) for exonuclease(s). Bi-directional resection releases SPO11-oligo complexes and exposes long ssDNA tails. ii, Two-step resection model involving MRE11 (green Pacman) handing off to other exonucle-olytic enzymes (black Pacman). iii, Sequencing adap-tors are ligated to duplex ends after removal of ssDNA. (B) Strategy for conditional deletion of *Mre11*. (C) Reduced MRN protein levels in *Mre11-cKO* mice. Immunoblotting is shown of whole testis lysates from 5–7 wk old animals. Numbers below the MRE11 blot indicate amounts of a dilution series of wild-type (wt) lysate or band intensities for *Mre11-cKO* and *Mre11-cHet* relative to wild type and normalized to the loading control (tubulin). (D) Representative PFA-fixed seminiferous tubule sections at 5–7 wk stained for MRE11 (brown color). In wild type, all cells except elongated spermatids stained positive for MRE11. In *Mre11-cKO*, tubules were categorized based on the presence of MRE11-positive spermatocytes. Arrowheads mark examples of MRE11-positive spermatogonia (red) or spermatocytes with (purple) or without (green) MRE11 staining. Scale bars, 50 μm. (E) Quantification of seminiferous tubule classes illustrated in panel D. Number of animals tested: n=1 for wild type and n=2 for *Mre11-cKO* (mean ± range). (F) Strand-specific S1-seq [reads per million mapped reads (RPM)] at a representative DSB hotspot. Signals from three (wild type) or two (*Spo11–/–* and *Mre11-cKO*) biological replicate libraries were averaged and plotted. The semi-synchronous first wave of spermatogenesis in juvenile mice enhances the signal:noise ratio in S1-seq libraries from wild type, but older mice (≥ 5 wk) are analyzed for *Mre11-cKO* because of their more penetrant phenotype (see Methods). Data are smoothed with a 151-bp Hann window. SPO11-oligo sequencing data here (top plot) and throughout are from Lange et al. (2016). The baseline of the y-axis for each plot is 0. (G) Stereotyped distribution of resection endpoints around DSB hotspots. Heatmaps (data in 40-bp bins) show strand-specific reads around SPO11-dependent DSB hotspots from wild type [n = 13,960 from Lange et al. (2016)]. Each hotspot is shown as a horizontal line, strongest at the top. Sequencing signals were locally normalized by dividing by the total signal in a 4001-bp window around each hotspot’s center. Each hotspot thus has a total value of 1, to facilitate comparisons of spatial patterns between hotspots of different strengths.

The most detailed understanding of meiotic resection is currently for budding yeast, which uses a two-step DSB-processing mechanism (**Figure 1A**)^3^. In the first step, the conserved Mre11-Rad50-Xrs2 (MRX) complex together with Sae2 (homologous to MRE11-RAD50-NBS1 (MRN) and CtIP in mouse) nicks the Spo11-bound strands using the endonuclease activity of Mre11 and degrades ssDNA towards the DSB using the 3’→5’ exonuclease activity of Mre11. We will refer to this as resection initiation or short-range resection. In the second step, the more processive 5’→3’ exonuclease activity of Exo1 further degrades ssDNA away from the DSB. We will refer to this step as long-range resection, although it should be noted that this is considerably shorter than the long-range resection that can occur in non-meiotic contexts^3^.

The combined action of these nucleases generates long 3’-terminal ssDNA and releases Spo11 still covalently bound to short oligonucleotides (oligos)^2,4-13^. The average resection length in wild-type yeast is ~800 nt and this is reduced by more than half (to ~375 nt) in *exo1* nuclease-deficient mutants^11,12^.

It is comparatively elusive how resection is executed in mammalian meiocytes. Similar to yeast, mouse DSB processing generates long 3’-terminal ssDNA (averaging ~1100 nt) and releases covalent SPO11-oligo complexes^7,14,15^. Unlike yeast, however, net resection length is decreased by only ~10% in mouse *Exo1* mutants, suggesting that EXO1 plays only a modest role in resection or is redundant with another nuclease(s)^14,15^.

The contribution of MRN to resection has been difficult to address, in part because of experimental challenges due to the embryonic lethality of knockout mouse models lacking MRN subunits^16-18^. Conditional deletion of *Nbs1* in germ cells resulted in reduced numbers of foci of RAD51 and other recombination proteins; this was interpreted to reflect a role for MRN in DSB processing, but resection per se was not directly assessed^19^. More recently, conditional *Rad50* deletion was found to cause an increased occurrence of unresected DSBs and to reduce resection tract lengths^20^. However, how MRN promotes resection remains unclear. Finally, it also remains unclear how resection length ties in with successful execution of meiotic recombination and thus with fertility.

We address these questions here through genome-wide analysis of meiotic DSB resection in mouse spermatocytes conditionally depleted of MRE11 or carrying various MRN mutations, including several that model hypomorphic genetic variants found in human chromosomal instability syndromes^21-28^. Our findings indicate that MRN is essential for resection initiation in mammals, but also, surprisingly, for most long-range resection. Our findings further reveal how resection defects alter the binding of recombination factors to ssDNA and impinge on faithful execution of meiosis.

## RESULTS

### MRE11 is required to initiate resection

To circumvent the lethality of an *Mre11* null mutation^17^, we used a conditional deletion strategy combining a floxed *Mre11* allele^29^ with *Ngn3-Cre*^30-32^ (**Figure 1B**). *Ngn3-Cre* is expressed in spermatogonia during the mitotic divisions prior to meiotic entry, but not in spermatogonial stem cells (SSCs, **Figure S1A**)^31,33^. We will refer to *Mre11*^*flox/–*^ *Ngn3-Cre*^*+*^ as *Mre11-cKO* (for conditional knockout), *Mre11*^*flox/+*^ *Ngn3-Cre*^*–*^ or *Mre11*^*flox/flox*^ *Ngn3-Cre*^*–*^ as phenotypic wild type (wt), and *Mre11*^*flox/+*^ *Ngn3-Cre*^*+*^ and *Mre11*^*flox/–*^ *Ngn3-Cre*^*–*^ as *cHet* and *Het*, respectively.

MRE11 protein was reduced by about 20 fold in immunoblots of whole testis extracts from young adult *Mre11-cKO* males (5 wk old, **Figure 1C**). We used immunohistochemistry on seminiferous tubule sections to determine which cell types retained MRE11 protein (**Figures 1D,E**). Virtually all cells were positive for MRE11 in wild type, including the layers of spermatogenic cells and Sertoli cells within the tubules as well as interstitial somatic cells.

In *Mre11-cKO* mice, most tubules contained a basal layer(s) of spermatogonia and spermatocytes but lacked later-stage germ cells, and 16.1% of tubules lacked spermatocytes entirely (detailed further below) (**Figures 1D,E**). Spermatogonia and interstitial cells remained uniformly MRE11-positive as expected, but spermatocytes, when present, either lacked detectable MRE11 or were a mix of MRE11-positive and -negative in the same tubule. These findings indicate that the *Mre11-flox* allele is recombined efficiently but incompletely by *Ngn3-Cre* expression in spermatogonia, resulting in a substantial population of spermatocytes entering meiosis with no or low MRE11 protein. As described before^29^, RAD50 and NBS1 levels were also decreased when MRE11 was depleted (**Figure 1C**).

We visualized resection endpoints genome-wide using S1-sequencing (S1-seq), in which genomic DNA is digested with the ssDNA-specific nuclease S1 and the blunted DNA ends are captured and sequenced (**Figure 1A**)^11,15,34^. For some experiments, we used a combination of exonuclease VII and exonuclease T from *Escherichia coli* instead of nuclease S1 (Exo7/T-seq; based on END-seq^14,35,36^).

Most DSBs are formed in hotspots whose locations can be identified by sequencing of SPO11 oligos^37,38^ (**Figure 1F**, top). S1-seq and Exo7/T-seq yield *Spo11*-dependent reads distributed ~0.3–2 kb away from hotspot centers with defined polarity: top-strand reads from right-ward-moving resection tracts and bottom-strand reads from leftward tracts^15^ (**Figures 1A,F,G and S1B,C**). Both methods also generate reads close to hotspot centers (**Figures 1F,G and S1B,C**). This central signal is thought to arise from recombination intermediates, possibly D loops^14,15^ (**Figure S1D**). Importantly, both S1-seq and Exo7/T-seq can also detect DSBs that have not been resected at all (**Figure S1E**)^11,14,15,39,40^.

S1-seq and Exo7/T-seq profiles at hotspots were dramatically altered in young adult *Mre11-cKO* mice (**Figures 1F,G and S1B,C**). Virtually all signals accumulated at hotspot centers in a pattern strikingly similar to SPO11 oligos in wild type, with a strong central cluster and weaker flanking clusters to either side (**Figures 1F,G and S1B,C**). These findings are markedly different from those in a recent *Rad50* conditional knockout^20^. We conclude that DSBs remain unresected in the absence of MRE11.

### MRE11 controls DSB numbers and restrains double cutting

To summarize global patterns and to facilitate comparisons between genotypes, we plotted genome-wide profiles around DSB hotspots by co-orienting and averaging the sequencing signal from top and bottom strand reads (**Figure 2A**). Compared to wild type, *Mre11-cKO* mice had elevated read counts at hotspots, with an increase well beyond the sample-to-sample variation in wild type (**Figures 2A,B**). In response to DSBs, MRN complexes activate ataxia-telangiectasia mutated (ATM) protein kinase (Tel1 in budding yeast)^41-43^. During meiosis, ATM/Tel1 regulates the number of DSBs via a negative feedback mechanism (reviewed in^44^) (**Figure 2C**). We therefore surmised that the increased sequencing read count in *Mre11-cKO* mice reflects, at least in part, an increase in the number of DSBs per cell because of a loss of MRN-dependent ATM activation.

**Figure 2.**
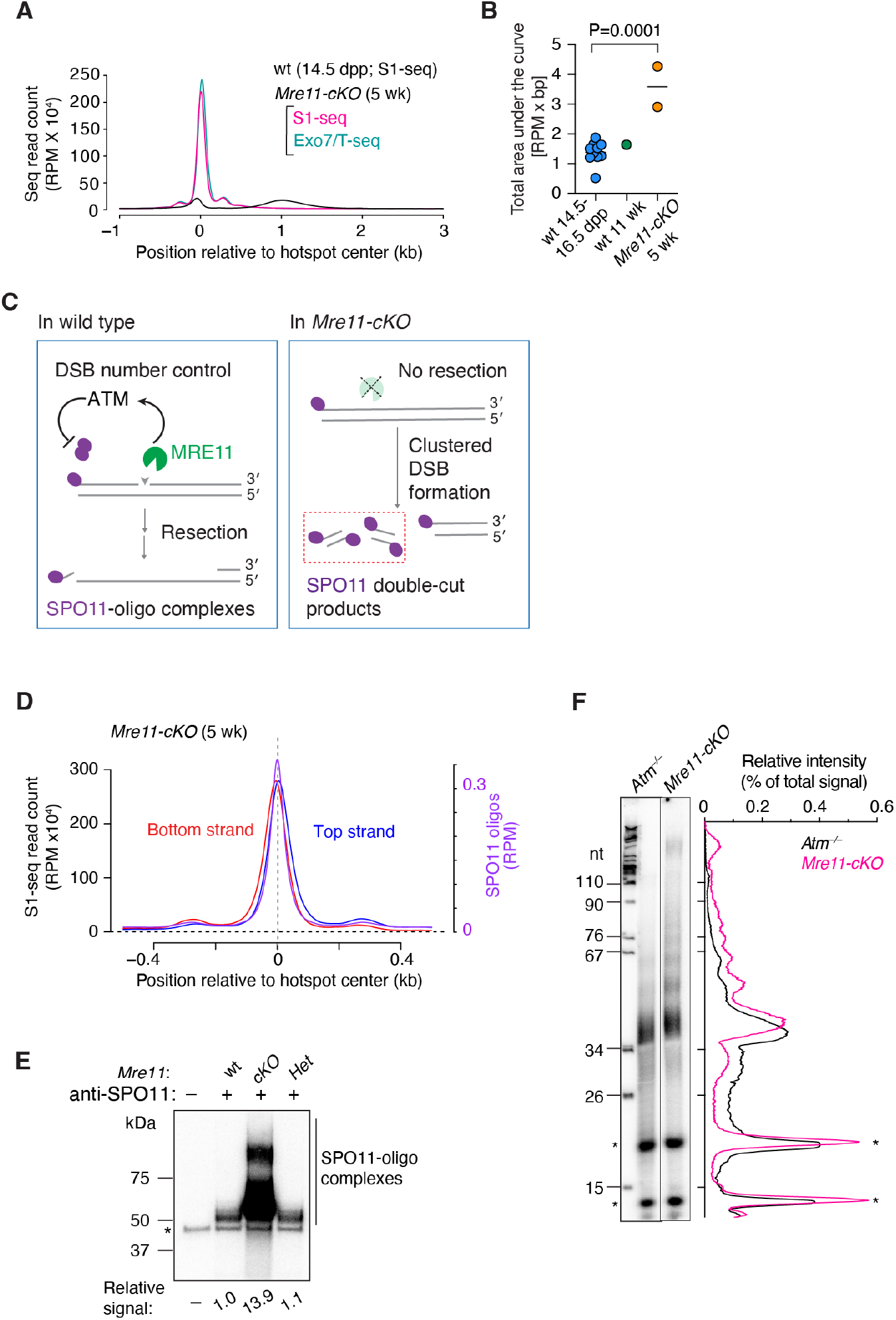
MRE11 fosters DSB number control. (A) Genome-average S1-seq and Exo7/T-seq at hotspots in 5-wk-old *Mre11-cKO* animals. The bottom-strand reads were flipped and combined with the top-strand reads and averaged (average of two biological replicates). Data are smoothed with a 151-bp Hann window. S1-seq in 14.5-dpp wild type is shown for comparison on the same scale (average of three biological replicates). (B) Comparison of S1-seq signal strengths for *Mre11-cKO* (n = 2) versus wild type (n = 10 for juvenile, 1 for adult). Each point is the area under the genome-average curve (as in panel A) from –1500 to +2500 bp relative to the hotspot center for an individual S1-seq replicate. The P value is from a two-tailed Student’s t test. (C) Role of MRE11 in ATM-mediated control of DSB formation and double cutting. Left, schematic illustrating ATM activation by MRE11 and subsequent negative feedback regulation of SPO11 activity. Right, schematic showing absence of resection and increase in SPO11 double cuts in the absence of MRE11. (D) Top- and bottom-strand reads at hotspot centers are offset in *Mre11-cKO*, consistent with double cutting. The plot shows the genome-average strand-specific profile close to hotspot centers, smoothed with a 51-bp Hann window. The average SPO11-oligo profile is shown for comparison. (E) Increased amounts of SPO11-oligo complexes with altered electrophoretic mobility in *Mre11-cKO*. A representative autoradiograph is shown of SPO11-oligo complexes immunoprecipitated from testis extracts from 5–6 wk old mice, radiolabeled with terminal transferase and [a-32P]-dCTP, and separated by SDS-PAGE. The signal intensity relative to wild type is indicated below. Asterisk: non-specific labeling artifact. (F) SPO11 oligo lengths in *Mre11-cKO*. SPO11 oligos were purified from two mice of the indicated genotypes by immunoprecipitation and protease digestion, then radiolabeled with terminal transferase and [a-32P]-GTP and separated by denaturing urea PAGE. Both samples were run on the same gel, but the intensity of the *Mre11-cKO* signal was increased approximately threefold to facilitate comparison. Asterisks: non-specific species; nt: marker lengths in nucleotides. Lane traces (background-subtracted and normalized to the total lane signal) are shown to the right.

If so, we anticipated that *Mre11-cKO* mutants would also display another hallmark of ATM deficiency, namely, a high frequency of double cutting in which multiple SPO11 complexes cut the same DNA molecule in close proximity^45-47^. In support, the top-strand signal of the central S1-seq peak was shifted to the right of the SPO11-oligo peak, and the bottom-strand signal was shifted to the left (**Figure 2D**). This pattern is distinct from the central signal derived from recombination intermediates in wild type (**Figure S2A**), but is as expected if multiple DSBs are formed on the same DNA molecules within hotspots because only the outermost DSB ends will be represented in the final sequencing library (**Figures 2C and S1E**). Double cutting may also explain why, unlike for SPO11 oligos in wild type, the subsidiary peak to the right of the central peak is stronger on average than the left-side subsidiary peak for top-strand reads, and vice versa for bottom-strand reads (**Figures 1G and 2D**).

We further tested for double cutting by radiolabeling the 3’ ends of SPO11-oligo complexes immunoprecipitated from whole-testis extracts. This assay does not detect SPO11 if it is still bound to the high molecular weight DNA of unresected DSBs, but double cutting can generate SPO11-oligo complexes in the absence of MRE11 nuclease activity^48^, as has been shown in budding yeast^45,47^ (**Figure 2C**). We indeed observed SPO11-oligo complexes in *Mre11-cKO* mice, and these were at levels more than ten-fold greater on a per-testis basis than in wild-type or heterozygote controls (**Figures 2E and S2B**), comparable to *Atm*^*–/–*^ mice^48^.

The major band of SPO11-oligo complexes from *Mre11-cKO* had slower mobility on SDS-PAGE than controls (**Figure 2E**). We therefore determined the size distribution of radiolabeled SPO11 oligos on denaturing PAGE (**Figure 2F**). Similar to SPO11 oligos purified from *Atm*^*–/–*^ mice compared side by side, those from *Mre11-cKO* mice displayed a major species migrating slightly slower than the 34-nt marker plus a ladder of slower migrating bands with ~10-nt periodicity. This periodicity is a diagnostic feature of SPO11 double cuts and may reflect geometric constraints on the orientation of adjacent SPO11 dimers relative to the DNA^45,47,49^. Interestingly, however, compared with *Atm*^*–/–*^, the *Mre11-cKO* sample displayed substantially less labeled material migrating faster than the 34-nt marker (**Figure 2F**), which is the position where SPO11 oligos released by resection are expected to run^48^. This difference reflects that *Atm*^*–/–*^ mice retain the ability to resect meiotic DSBs^14,15^ but *Mre11-cKO* mice do not.

We also examined resection patterns in juvenile (14.5 dpp) *Mre11-cKO* mice. Immunoblotting of whole-testis extracts showed that MRE11 protein was reduced, but to a lesser degree than in adults (~twofold lower than controls; **Figure S2C**). Similar to adults, these mice showed a greatly elevated central S1-seq signal at hotspots that had the spatial patterns diagnostic of unresected DSBs and double cutting (**Figures S2D–F**). Unlike adults, however, juveniles also had a substantial fraction of resected DSBs, although with a resection length distribution that was markedly shorter (modal length 897 nt) than in wild type (modal length 1015 nt) (**Figure S2F**). It is formally possible that spermatocytes in the initial wave of meiosis in juveniles have an additional (MRE11-independent) resection initiation mechanism that is not available in adults. We consider it more likely, however, that the weaker phenotype in juveniles reflects differences in the timing of Cre-mediated excision relative to DSB formation and/or differences in the kinetics of MRE11 protein rundown after excision.

### Spermatogenesis failure in *Mre11*-deficient mice

Adult *Mre11-cKO* mice had significantly smaller testes than littermate wild-type or heterozygous controls (**Figures 3A,B**). In seminiferous tubule sections, layers of spermatogonia and spermatocytes were present but postmeiotic cells were largely absent, sections of epididymis appeared empty, and epididymal sperm counts were greatly reduced (**Figures 3A,C**). All *Mre11-cKO* tubules contained one or more layers of cells that stained positive for the mouse VASA homolog DDX4 (**Figure S3A**), which is expressed from spermatogonia to the round spermatid stage^50^. Tubules with apoptotic cells were also increased in frequency (**Figures S3B,C**). These *Mre11-cKO* phenotypes remained consistent to at least 16 wk of age (the oldest tested) (**Figures S3D**). These results, together with MRE11 staining (**Figure 1D**), support the interpretation that *Mre11-cKO* mice stably maintain an MRE11-positive SSC population that can initiate multiple waves of spermatogenesis, but most spermatocytes in each wave are then eliminated by apoptosis. We conclude that MRE11 protein is essential for male fertility in mice, as are NBS1 and RAD50^19,20^.

**Figure 3.**
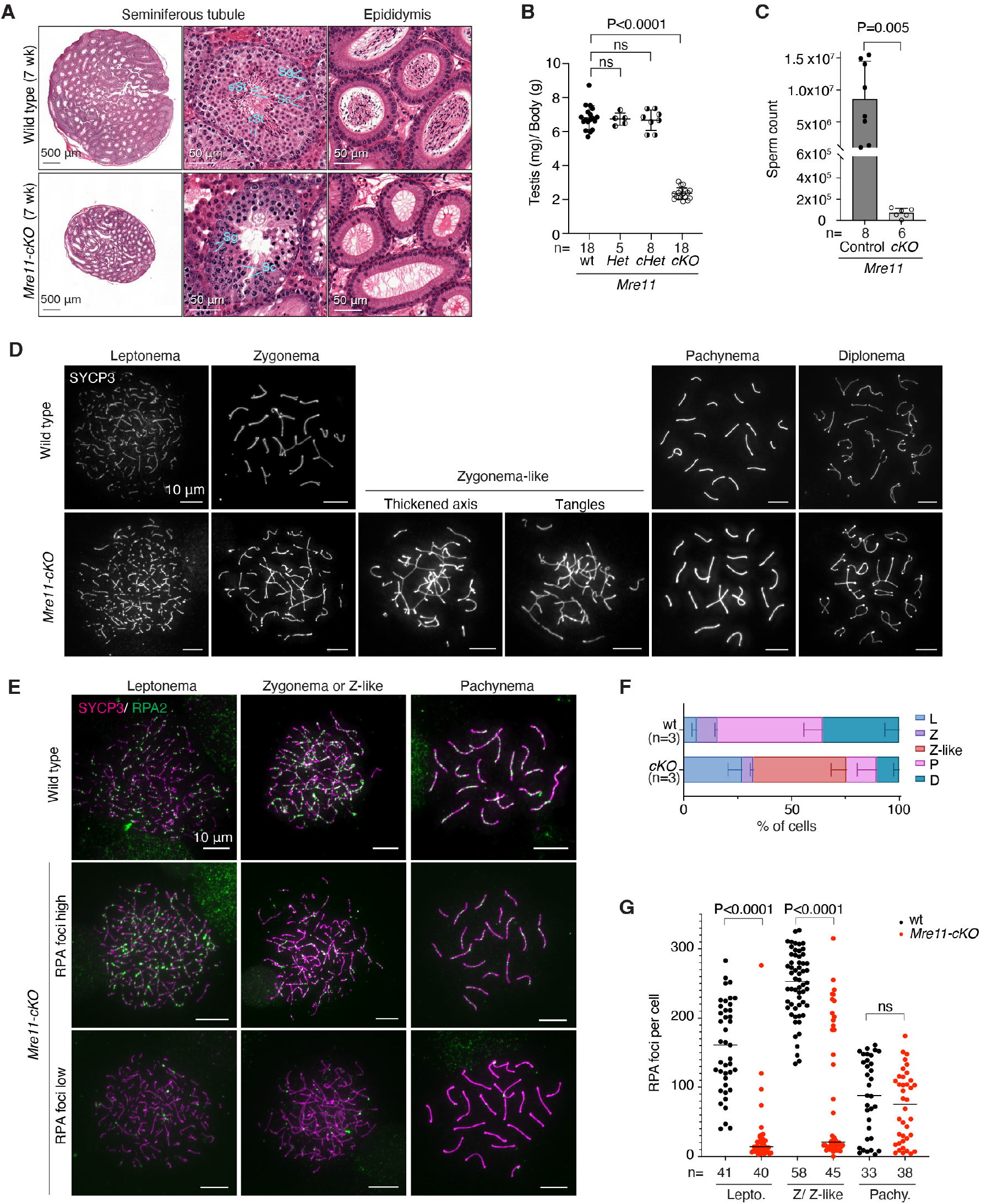
Spermatogenesis defects in *Mre11*-deficient mice. (A) Bouin’s fixed testis and epididymis sections stained with hematoxylin and eosin (H&E). Sg: spermatogonia, Sc: spermatocyte, rSt: round spermatid, eSt: elongated spermatid. (B) Ratios of testis weight (mg per pair) to body weight (g). Each point represents one animal; error bars indicate mean ± SD. The results of Student’s t tests are shown; ns, not significant (P > 0.05). (C) Sperm counts (mean ± SD; P value from Student’s t test). (D) Representative spreads of wild-type or *Mre11-cKO* spermatocytes showing normal SYCP3 staining at the indicated stages or abnormal (“Zygonema-like”) cells with thickened and/or tangled axes. (E) Representative RPA2 staining on spermatocyte chromosome spreads. Examples are shown of *Mre11-cKO* cells with either low or high RPA2 focus counts. (F) Spermatocyte stages based on SYCP3 staining (L: leptonema, Z: zygonema, Z-like: zygonema-like, P: pachynema, D: diplonema). Error bars indicate mean ± SD from three mice of each genotype. (G) RPA2 focus numbers. Error bars represent mean ± SD for the indicated number of cells counted from 2 mice of each genotype. The stage of each cell was determined from the SYCP3 signal. The results of two-tailed Mann-Whitney U tests are shown.

To monitor chromosome dynamics and progression through prophase I, we immunostained spermatocyte chromosome spreads for the axis protein SYCP3, with or without costaining for the RPA2 subunit of the ssDNA-binding protein RPA. In wild type, SYCP3 forms short lines in leptonema as axes begin to form; the axes elongate and synapse with homologous partners to form stretches of synaptonemal complex (SC) in zygonema; SC spans the length of each pair of autosomes in pachynema; and then SC disassembles in diplonema (**Figures 3D and S3E**). RPA2 forms large numbers of foci on resected DSBs in leptonema and zygonema, then these foci decrease in number over the course of pachynema as DSB repair is completed (**Figure 3E**)^51^.

Spermatocytes in *Mre11-cKO* mice deviated substantially from the normal patterns. We observed cells with short SYCP3 axes and no SC, similar to normal leptonema but representing a higher fraction of total spermatocytes than in wild type (**Figures 3D,F and S3E**). Most cells had very few RPA2 foci (**Figures 3E,G**). We also observed cells with axes of various lengths along with SC, but most of these differed from normal zygotene cells in having only short stretches of staining for SC central region component SYCP1 and/or tangles of axes (**Figure S3E**). These “zygotene-like” cells were the most abundant class in *Mre11-cKO* (**Figure 3F**) and most were again highly depleted for RPA2 foci compared to wild type (**Figures 3E,G**). The deficit of RPA2 foci corroborates the resection defect seen in S1-seq. These results also show, not surprisingly, that unresected DSBs cannot support homologous pairing and synapsis.

In addition to these aberrant cells, there were also morphologically normal pachytene and diplotene cells, but much fewer than in wild type (**Figures 3D,F and S3E**). RPA2 focus numbers in the pachytene cells were indistinguishable from wild type (**Figure 3G**). We also observed a small fraction of leptotene and zygotene cells that had RPA2 focus numbers in the normal range for these stages (**Figures 3E,G**). We infer that these more normal-looking cells correspond to the MRE11-positive subpopulation of spermatocytes (**Figure 1D**), perhaps cells that escaped Cre-mediated excision. If so, these escapers appear to be too small a fraction of the total leptotene and zygotene cells (**Figure 3F**) to contribute an appreciable resection signal in population-average sequencing maps (**Figures 1F,G and 2A**).

### Nuclease-dead MRE11 reveals an unexpected role in long-range resection

To test whether MRE11 nuclease activity is required for meiotic resection, we examined mice with the invariant active-site residue His-129 substituted to asparagine^29^. Human MRN complexes with this mutation lack nuclease activity in vitro^52,53^. *Mre11-H129N* is cell-lethal in chicken and human cells^54^ and is embryonically lethal in mice^29^, but the phenotype in mammalian meiosis has not been addressed. To circumvent the lethality, we generated *Mre11*^*H129N/flox*^ mice carrying *Ngn3-Cre* (hereafter *Mre11-cHN*) (**Figure 4A**). Total MRE11 protein levels were normal in testis extracts from *Mre11-cHN* mice (**Figure 4B**), consistent with previous demonstration that the mutation does not destabilize the protein^29^.

**Figure 4.**
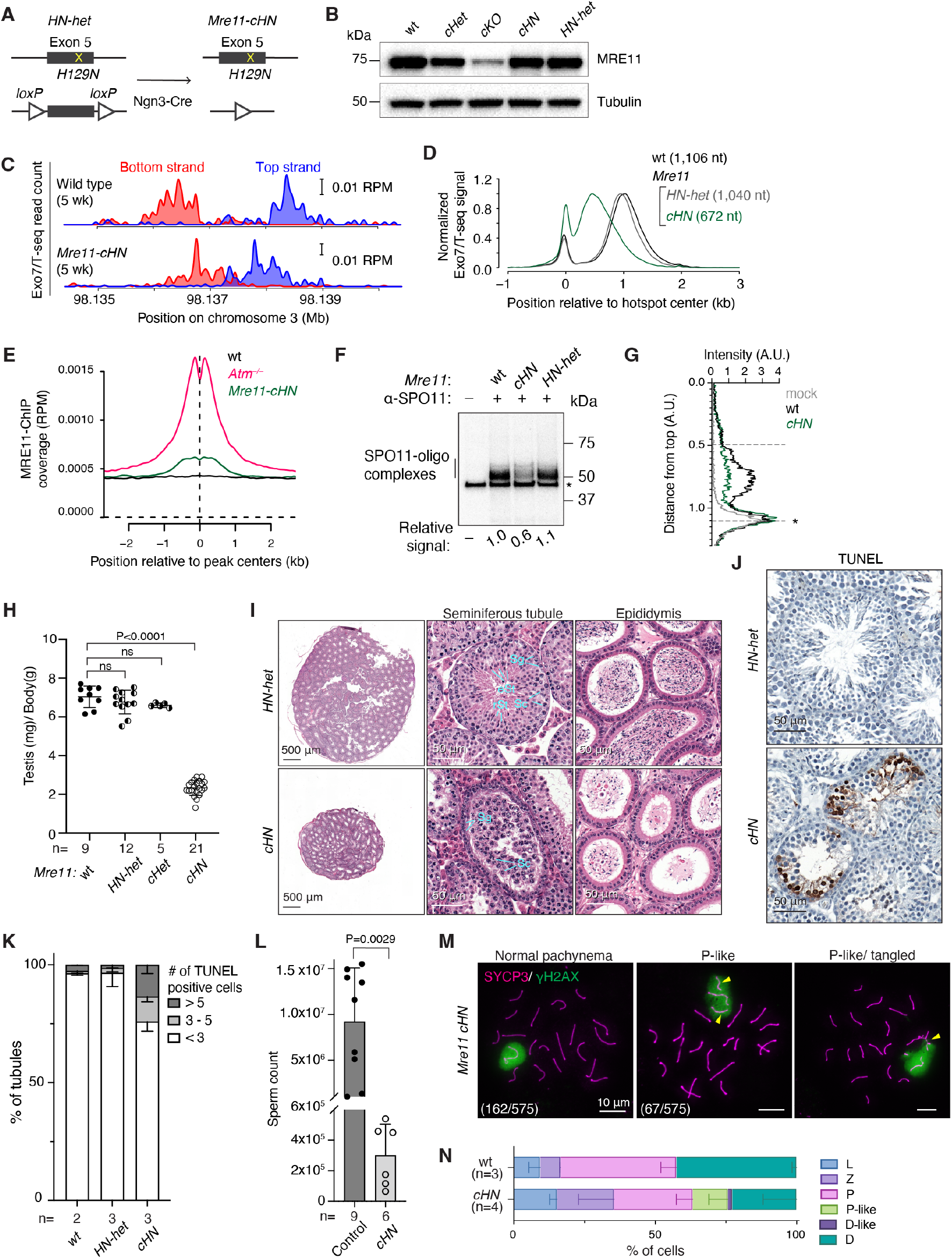
Defects in long-range resection and spermatogenesis in mice expressing nuclease-dead MRE11. (A) Conditional *Mre11-H129N* strategy combining *Ngn3-Cre* with compound heterozygous floxed and missense *Mre11* exon 5 alleles. (B) Total MRE11 protein levels are normal in *Mre11-cHN* mice. Immunoblots are shown of whole testis lysates from 5–7 wk old mice. In this experiment, the phenotypically wild-type control (wt) was *Mre11wt/flox Ngn3-Cre–* and the heterozygous HN strain (*HN-Het*) was *Mre11H129N/flox Ngn3-Cre–*. (C) Exo7/T-seq at the same hotspot shown in **Figure 1F**. (D) Genome-average Exo7/T-seq patterns at hotspots in 5-wk-old animals. The sequencing signal from top and bottom strand reads was co-oriented, averaged, and smoothed with a 151-bp Hann window, then internally normalized by setting the height of the resection peak to a value of 1. The normalization facilitates comparison of the shapes of the resection profiles separate from sample-to-sample variation in the signal:noise ratio of the sequencing read frequencies34. Non-normalized plots are shown in **Figure S4C**. Two biological replicates each for wild type and *Mre11-cHN* were averaged and plotted. The profile for *HN-Het* was from a single library. (E) MRE11 accumulation at hotspots in *Mre11-cHN*. MRE11 ChIP-seq signal was averaged around hotspots for *Mre11-cHN* (n=3 biological replicates), wild type (n=2), and *Atm–/–* (n=1). (F,G) Reduced amount of SPO11-oligo complexes in *Mre11-cHN*. An autoradiograph of radiolabeled complexes immunoprecipitated from testis extracts (F) and lane profiles (G) are shown. (H) Reduced testis sizes. Error bars indicate mean ± SD. Results of Student’s t tests are shown. (I) Bouin’s fixed and H&E-stained seminiferous tubule and epididymis sections (7 wk old; cell type labels as in **Figure 3A**). (J,K) Increased spermatocyte apoptosis in *Mre11-cHN* mice (7 wk). Representative TUNEL staining of seminiferous tubule sections (J) and quantification (K) are shown. Error bars in K indicate mean ± range. Wild type is reproduced from **Figure S3C**. (L) Sperm counts (mean ± SD; P value from a Student’s t test). The control is a combination of wild type, *HN-het*, and *cHet* animals. (M) Representative spreads of wild-type or *Mre11-cHN* spermatocytes showing either normally synapsed SYCP3 staining of autosomes and normal appearing γH2AX-positive sex body (left) or pachytene-like cells with unpaired or tangled autosomes (middle and right micrographs), which were associated with the γH2AX-positive domain (arrowheads). At lower left, the number of pachytene or pachytene-like cells observed and total number of SYCP3-positive cells counted. (N) Spermatocyte stages based on SYCP3 and γH2AX staining. Error bars indicate mean ± SD.

Because the equivalent budding and fission yeast mutants form meiotic DSBs but cannot resect them^6,55^, we expected that *Mre11-cHN* mice would accumulate only unresected DSBs if the wild-type MRE11 protein expressed from the floxed allele were sufficiently depleted (**Figure S4A**, left). On the other hand, if some wild-type MRE11 protein persisted long enough to support resection initiation at a subset of DSBs in each cell, or at all DSBs in a subset of escapee cells, we instead expected to observe that some or all of the DSBs would be fully resected (**Figure S4A**, right). The latter prediction was premised on the idea that the long-range nuclease(s) should be able to complete resection normally as long as MRE11 endonuclease had incised the DNA.

Surprisingly, however, Exo7/T-seq patterns in young adult mice matched neither of these predictions when considering either individual hotspots (**Figures 4C and S4B**) or genome-average profiles (**Figures 4D and S4C**). We note two main findings. First, there was a modest increase in the relative amount of the sequencing signal at hotspot centers (**Figure 4D**), suggesting that some unresected DSBs are present. Consistent with this interpretation, we also observed a slight increase in the sequencing signal at hotspot centers on the nonhomologous parts of the X and Y chromosomes (**Figure S4D**). Central S1-seq or Exo7/T-seq signal from recombination intermediates does not appear at X and Y hotspots in wild type, but it can be detected when there are unresected DSBs, such as in *Atm*^*–/–*^ mutants^14,15^ or *Mre11-cKO* (**Figure S4E**).

To further test for unresected DSBs, we performed chromatin immunoprecipitation (ChIP-seq) for MRE11, which was previously demonstrated to accumulate at hotspots when unresected DSBs are present in *Atm*^*–/–*^ mutants, but not in wild type where all DSBs are resected^14^. Indeed, we observed a clear MRE11 ChIP-seq signal at hotspots in *Mre11-cHN* mice, albeit substantially less than in *Atm*^*–/–*^ (**Figures 4E and S4F,G**). Taken together, these results suggest that a small fraction of DSBs remains unresected in *Mre11-cHN* mice. If so, however, this resection initiation defect is very weak compared to *Atm*^*–/–*^ and especially *Mre11-cKO*, perhaps reflecting insufficient elimination of wild-type MRE11 protein in this conditional model (see Discussion).

The second and more surprising finding was that resection tracts were substantially shorter than normal in *Mre11-cHN*, averaging only 672 nt (**Figure 4D**). This defect was reproducible and stable, remaining the same with increasing age up to 11 wk (the oldest age tested; **Figure S4H**). We also observed a slight resection defect in mice heterozygous for *Mre11-H129N* (*Mre11*^*H129N/wt*^ with *Ngn3-Cre*). Resection tracts in these mice averaged 1040 nt (94% of wild type) (**Figure 4D**), suggesting a weak dominant-negative effect of MRE11-H129N protein. Observing shorter average resection lengths reveals that MRE11 nuclease activity contributes to long-range resection, not just to resection initiation.

Unlike in *Atm*^*–/–*^ and *Mre11-cKO*, the amount of SPO11-oligo complexes was not increased in *Mre11-cHN* (**Figures 4F and S4I**), consistent with MRE11-HN protein being competent to activate ATM/ Tel1 in both yeast and mouse^29,53,56,57^. Instead, the yield on a per-testis basis was reduced by about half compared to wild type. The total Exo7/ T-seq signal recovered at hotspots was also diminished in *Mre11-cHN* (**Figure S4J**). These decreases may reflect reduced DSB frequency and/ or loss of signal because of spermatocyte apoptosis (described below). We also observed a shift in the migration of SPO11-oligo complexes on SDS-PAGE gels, with faster migrating species more depleted than slower migrating ones (**Figure 4G**). We posit that the presence of catalytically inactive MRE11 protein sometimes occludes potential nicking sites closer to SPO11, biasing the population of SPO11 oligos generated by the residual wild-type MRE11 protein to larger sizes than normal.

We were unable to assess a role for CtIP—a cofactor needed for MRN nuclease activity^3^ because conditional deletion of *Ctip* eliminated spermatogonia prior to meiotic entry (**Figures S4K–N**). We also tested *Rad50*^*S/S*^ mice but found normal resection (**Figure S4O**), likely because this *Rad50-K22M* mutation was based on a relatively weak yeast allele because mimics of stronger alleles are cell-lethal^23^. Thus, available mouse mutants are unable to definitively test the requirement for MRN nuclease in resection initiation.

### Spermatogenesis defects in mice with nuclease-dead MRE11

*Mre11-cHN* mice had pronounced defects in meiotic progression. Their testes were considerably smaller than those of control littermates (**Figure 4H**); seminiferous tubules contained spermatogonia and early stage spermatocytes but were depleted of spermatids and had an increased frequency of spermatocyte apoptosis (**Figures 4I–K**); and epididymal sperm counts were greatly diminished (**Figures 4I,L**). The rare residual sperm detected may have arisen from escapers in which Cre-mediated excision failed and/or from a subset of cells in which wild-type MRE11 protein persisted long enough after excision to support successful meiosis.

When prophase I stages were evaluated by staining spreads for SYCP3 and phosphorylated H2AX (γH2AX), we observed a subset of pachytene-like cells with unpaired or tangled chromosomes (**Figures 4M,N**), indicative of sporadic synaptic failure. We also observed an increase in the proportions of leptotene and zygotene cells relative to later stages (pachytene, pachytene-like, and diplotene) (**Figure 4N**). This increase may reflect a delay in completing synapsis, preferential apoptotic elimination of cells in later prophase I, or both.

### Reduced resection lengths in MRN hypomorphic mutants

We further tested for MRN roles in meiotic resection using mice homozygous for hypomorphic mutations modeled on human disease alleles^58^ (**Figure 5A**). *Mre11-ATLD1* is a nonsense mutation near the 3’ end of the coding sequence, recapitulating a mutation found in ataxia telangiectasia-like disorder^21^. *Nbs1ΔB*, which models Nijmegen breakage syndrome, produces an NBS1 protein lacking the N-terminal FHA and BRCT domains^26^. *Mre11-ATLD1* and *Nbs1ΔB* cause intra-S and G2/M checkpoint defects, DNA damage sensitivity, chromosomal instability, and reduced ATM activity in cultured cells and they cause subfertility in mice, particularly in females^21,26,59^. The mutations also reduce protein stability and MRN nuclease activity in vitro^60^, but they have not previously been linked to overt resection defects.

**Figure 5.**
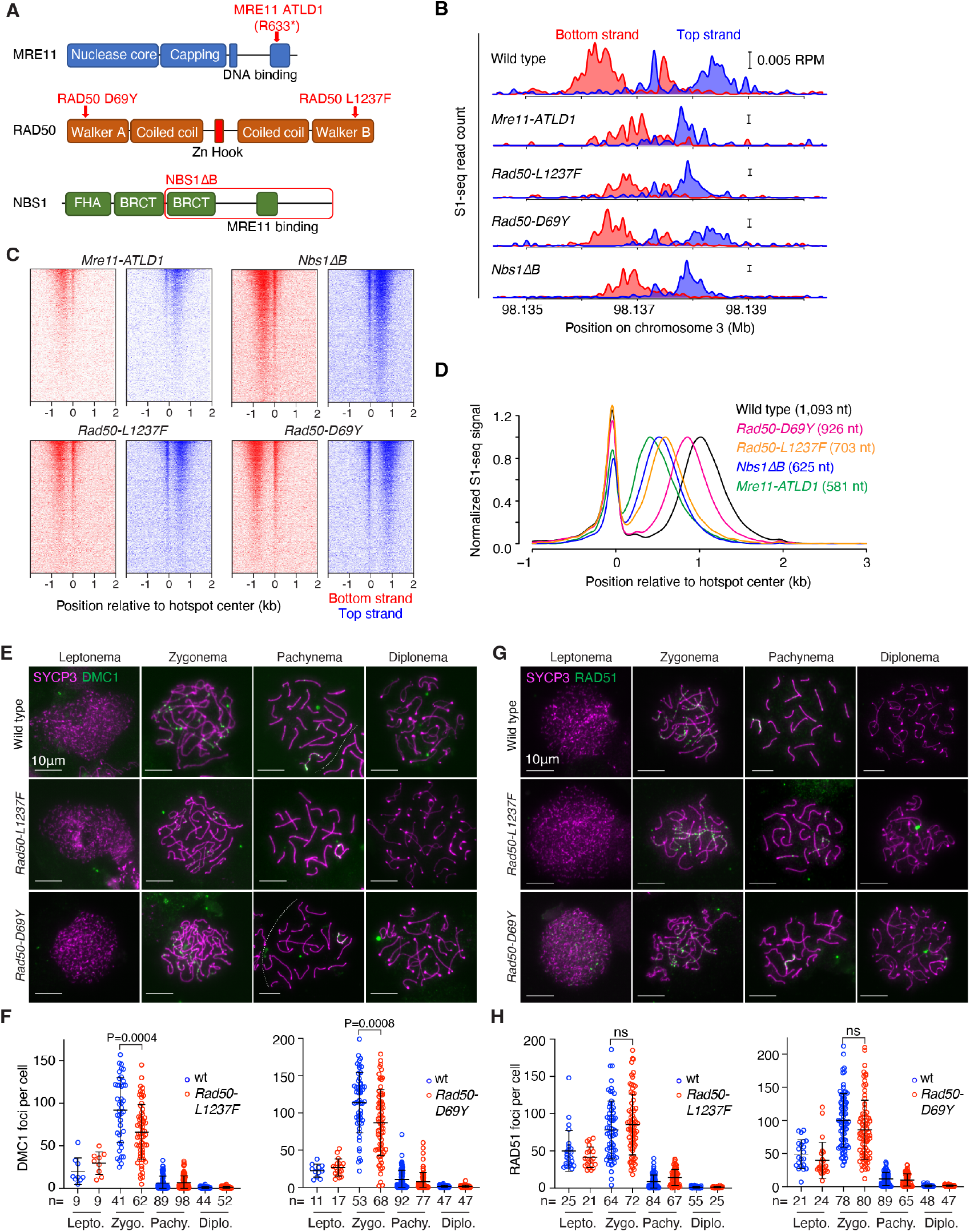
Reduced resection length in MRN hypomorphs. (A) Schematic of MRN hypomorphs. MRE11 R632^*^ (R632 → stop) mimics human R633^*^ identified in ATLD patients21. *Nbs1ΔB* produces an 80 kDa protein (red box), mimicking the *657del5* allele found in 95% of NBS patients26. (B) S1-seq at the hotspot from **Figure 1F**, from 14.5-dpp mice. Wild type is reproduced from **Figure 1F**. Two biological replicates were averaged for each of the other genotypes. (C) Heatmaps (data in 40-bp bins) of S1-seq reads around DSB hotspots. (D) Genome-average S1-seq at hotspots from 14.5-dpp mice. Wild type is reproduced from **Figure 2A**. Data are smoothed with a 151-bp Hann window and normalized as described in **Figure 4D**. (E–H) Minimal if any changes in RAD51 and DMC1 focus numbers during meiosis in *Rad50-L1237F* or *Rad50-D69Y* mice. Representative spreads stained for SYCP3 and DMC1 (E) and RAD51 (G) along with quantification (F, H) are shown. Each dot in the graphs is the focus count from a single nucleus; error bars indicate means ± SD. Results of unpaired t tests are shown for zygonema; differences at the other stages were not statistically significant (P > 0.05).

We also examined homozygous *Rad50-D69Y* and *Rad50-L1237F* mutants, which model recurrent mutations in human cancer affecting residues in the Walker A and B ATPase motifs, respectively^61,62^ (**Figure 5A**). Studies in *S. cerevisiae* and cultured mouse cells indicated that the human *RAD50-L1240F* mutant (*L1237F* in mouse) is a separation-of-function allele that is largely proficient in DNA repair, but defective in the activation of Tel1/ATM^61^. Similar outcomes were observed for modeling of human *RAD50-D69Y* in *S. cerevisiae*^62^.

*Rad50-L1237F* and *Rad50-D69Y* mice were derived by CRISPR-mediated mutagenesis. MEFs from *Rad50-D69Y* mice displayed reduced ionizing radiation (IR)-provoked phosphorylation of KAP1 (an ATM target^63^) (**Figure S5A**); elevated mitotic indices after IR, consistent with defects in the ATM-dependent G2/M checkpoint (**Figure S5B**); and hypersensitivity to camptothecin when ATR was inhibited, as expected if compromised ATM activation cannot compensate for loss of ATR activity (**Figure S5C**). These findings confirm the separation of function predicted from prior studies in *S. cerevisiae*^62^, and verify that *Rad50-D69Y*, like *Rad50-1237F*^61^, is defective for ATM activation.

S1-seq maps from juvenile males demonstrated resection defects in all of these MRN mutants, demonstrating a striking allelic series (**Figures 5B–D**). These findings reinforce the conclusion that the MRN complex must be fully functional to support normal long-range resection, i.e., that MRN roles in DSB processing are not limited just to resection initiation. There was only a slight increase in central S1-seq signal at hotspot centers on the sex chromosomes in *Nbs1ΔB*, suggestive of a small number of unresected DSBs, but little if any evidence for this in the other mutants (**Figure S5D**).

Spermatogenesis in *Nbs1ΔB* and *Mre11-ATLD1* mutant males was previously shown to be nearly normal, with only mildly delayed temporal progression and persistent RAD51 localization that was attributed, at least in part, to reduced functionality of these MRN hypomorphs in meiotic DSB repair^21,59^. The substantially shorter resection lengths we now observe in these mutants may contribute to this apparent DSB repair delay. Spermatogenesis also appeared to be largely normal in *Rad50-L1237F* and -*D69Y* mutants. Testis sizes and sperm counts were indistinguishable from littermate controls (**Figure S5E–G**); the frequency of spermatocyte apoptosis was not significantly increased (**Figure S5H**); the distribution of prophase I stages was not substantially altered (**Figure S5I**); and the appearance and disappearance of foci of strand exchange proteins occurred normally, with the exception that both mutants showed a statistically significant but quantitatively small reduction in the number of DMC1 foci at zygonema (**Figures 5E–H**). These findings show that meiotic recombination is robust in the face of even substantial deviations from normal resection lengths (e.g., up to the ~47% reduction in *Mre11-ATLD1*).

### Side-by-side spatial organization of RAD51 and DMC1 is maintained in *Nbs1ΔB* mice

We tested whether the shortened resection lengths in *Nbs1ΔB* mice altered the occupancy and spatial disposition of ssDNA binding proteins RPA, DMC1, and RAD51 by performing ChIP followed by ssDNA sequencing (SSDS)^64,65^. In wild type, DMC1 tends to bind closer to the broken end, RAD51 binds nearer to the ssDNA/dsDNA boundary, and RPA can be found essentially anywhere along the ssDNA^64^ (**Figure 6A**). The SSDS signal for all three proteins was more compressed toward hotspot centers in the *Nbs1ΔB* mutant, as expected from the decreased length of ssDNA available, but the spatial pattern of DSB-proximal DMC1 and DSB-distal RAD51 was maintained (**Figure 6B**).

**Figure 6.**
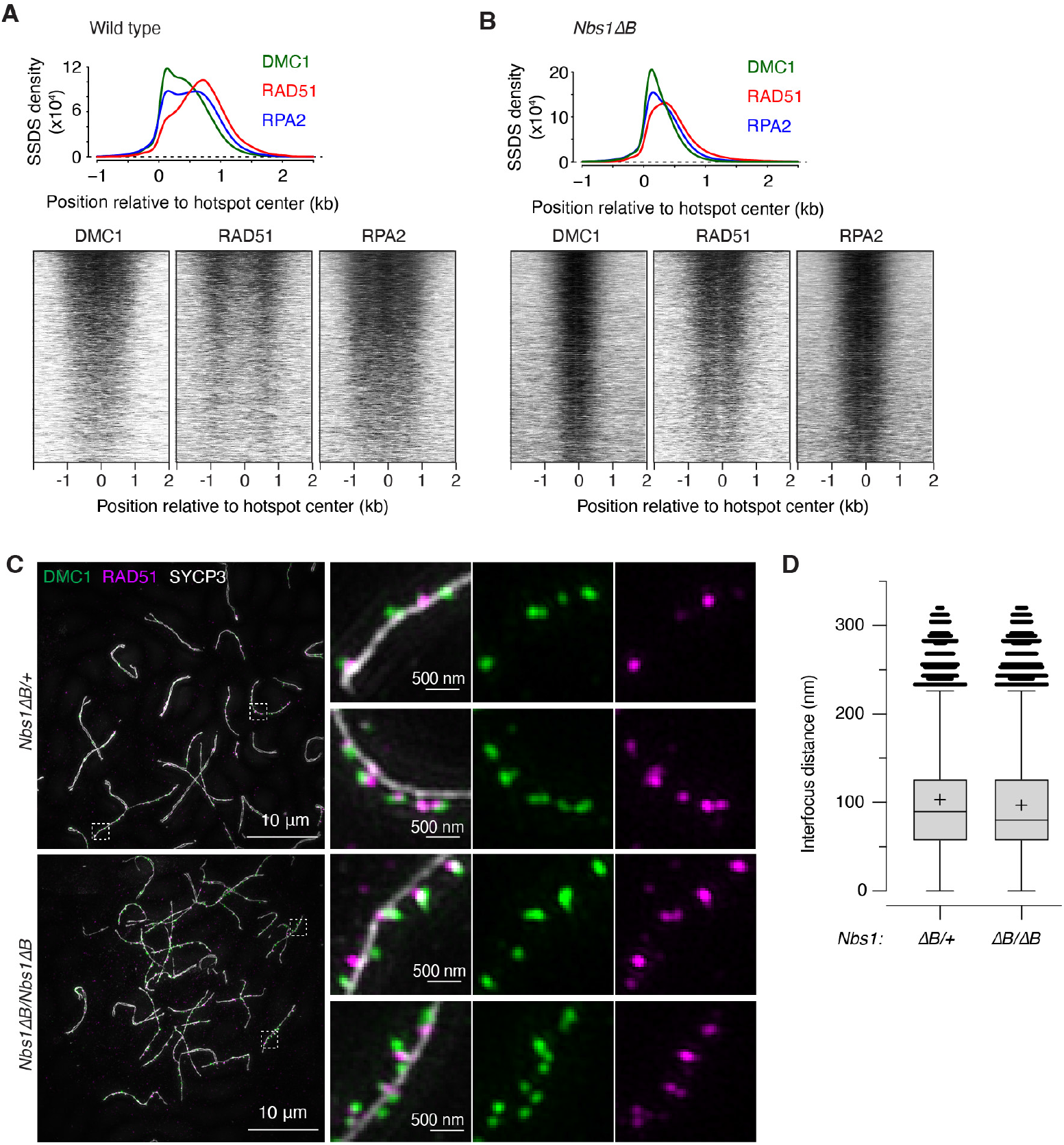
Side-by-side spatial organization of RAD51 and DMC1 is maintained despite shortened resection in *Nbs1ΔB* mice. (A, B) Genome-wide profiles of DMC1, RAD51, and RPA2 ChIP followed by SSDS in wild type (A) or *Nbs1ΔB* (B). In the graphs above, top- and bottom-strand reads around hotspots were co-oriented, combined and averaged. These maps represent sequencing coverage rather than endpoint counts reported for S1- or Exo7/T-seq. Data are smoothed with a 151-bp Hann window. An estimated background was removed by subtracting the value of signal 2.5 kb away from the hotspot center, then profiles were normalized by setting the area under each curve (from –1.0 to 2.5 kb) to 1. Heatmaps below show SSDS signals in 40-bp bins, locally normalized by dividing each signal by the total signal in a 4001-bp window around each hotspot’s center. Each hotspot thus has a total value of 1, so that spatial patterns can be compared between hotspots of different strengths. (C) Representative SIM images of spread spermatocytes from 14.5-dpp mice. Insets show zoomed views of the regions indicated by dashed boxes. (D) DMC1 to RAD51 distances for axis-associated co-foci. Tukey box plots are shown for DMC1 foci that were £450 nm from an axis and £320 nm from the nearest RAD51 focus. The plus signs indicate means; horizontal lines are medians; box edges are interquartile range; whiskers indicate the most extreme data points which are ≤1.5 times the interquartile range from the box; individual points are outliers.

Previous studies in yeast and mice showed that foci of RAD51 and DMC1 often appear side by side^64,66,67^. We tested using structured illumination microscopy (SIM) whether this organization at individual DSB sites was retained in *Nbs1ΔB* mice, comparing homozygous mutants with heterozygous littermate controls (which have nearly normal resection lengths (**Figure S6A**)). Similar to previous results^64^, a majority of the foci of both DMC1 (93.8%) and RAD51 (79.4%) were located close to chromosome axes in *Nbs1*^*ΔB/+*^ heterozygotes (**Figures 6C and S6B and Table S1**). Moreover, the foci often occurred together in pairs: 80.7% of axis-associated DMC1 foci and 77.4% of of axis-associated RAD51 foci were within 320 nm of the other protein (**Figures 6C and S6C and Table S1**). The high degrees of both axis association and colocalization were retained in *Nbs1*^*ΔB/ΔB*^ homozygotes (94.7% axis association for DMC1 and 86.3% for RAD51; 84.8% colocalization for DMC1 foci and 82.3% for RAD51) (**Figures 6C and S6B,C and Table S1**).

DMC1 and RAD51 foci were nearly always offset from one another,with a distance in the controls of 103.3 ± 67.5 nm (mean ± SD) between DMC1 foci and their RAD51 mates (**Figure 6D and Table S1**). This distance was decreased significantly in *Nbs1ΔB* mutants (P = 2.2 ′ 10^−16^, t test), but with a small effect size (96.8 nm ± 66.5).

Two conclusions emerge. First, the decreased distance between DMC1 and RAD51 foci when resection lengths are reduced supports the interpretation that paired foci represent binding of both proteins to the same ssDNA tail and not to separate DSB ends, similar to the situation inferred from yeast and mouse studies^66,67^. Second, the effect of the *Nbs1ΔB* mutation on interfocus distances (less than 7% decrease on average) was much smaller than the effect on both resection length (43% decrease, **Figure 5D**) and ChIP profiles (**Figure 6B**). This difference between cytology and population-average molecular assays shows that ssDNA length is not the most important determinant of the three-dimensional spatial organization of DSB repair foci. This conclusion in turn fits with the idea that individual DMC1 and RAD51 filaments are fairly short, estimated at ~100 nt in yeast and ~400 nt in mouse^66,67^. Because filaments are only long enough to occupy a small fraction of the available ssDNA, even a substantial reduction in resection lengths should not cause a proportionate decrease in the size of cytologically observable foci.

### Additive contributions of EXO1 and the NBS1 N terminus to long-range resection

In vitro, yeast MRX promotes Exo1 activity by providing nicks for Exo1 to act on and by fostering Exo1 binding to DNA ends^68-71^. To query the interplay between MRN and EXO1 in meiotic resection, we generated double mutant mice homozygous for both *Nbs1ΔB* and *Exo1* mutations. An *Exo1* null is synthetically lethal with *Nbs1ΔB*^72^, so we used the nuclease-deficient *Exo1-D173A (DA)* allele^73^, reasoning that the milder somatic phenotype of *Exo1DA*^74^ might allow us to obtain double mutants. Indeed, *Nbs1ΔB Exo1DA* doubly homozygous mice were viable, albeit recovered at a sub-mendelian frequency (**Figure S7A**) and exhibiting lower body weights (**Figure S7B**).

*Nbs1ΔB Exo1DA* double mutants showed a 111-nt decrease in the average resection length compared to *Nbs1ΔB* alone (**Figures 7A,B and S7C**). Since this is similar to the 126-nt average decrease in *Exo1DA* compared to wild type, EXO1 appears to work additively with MRN. We also conclude that the *Nbs1ΔB* resection defect is not attributable to a defect in EXO1 activity.

**Figure 7.**
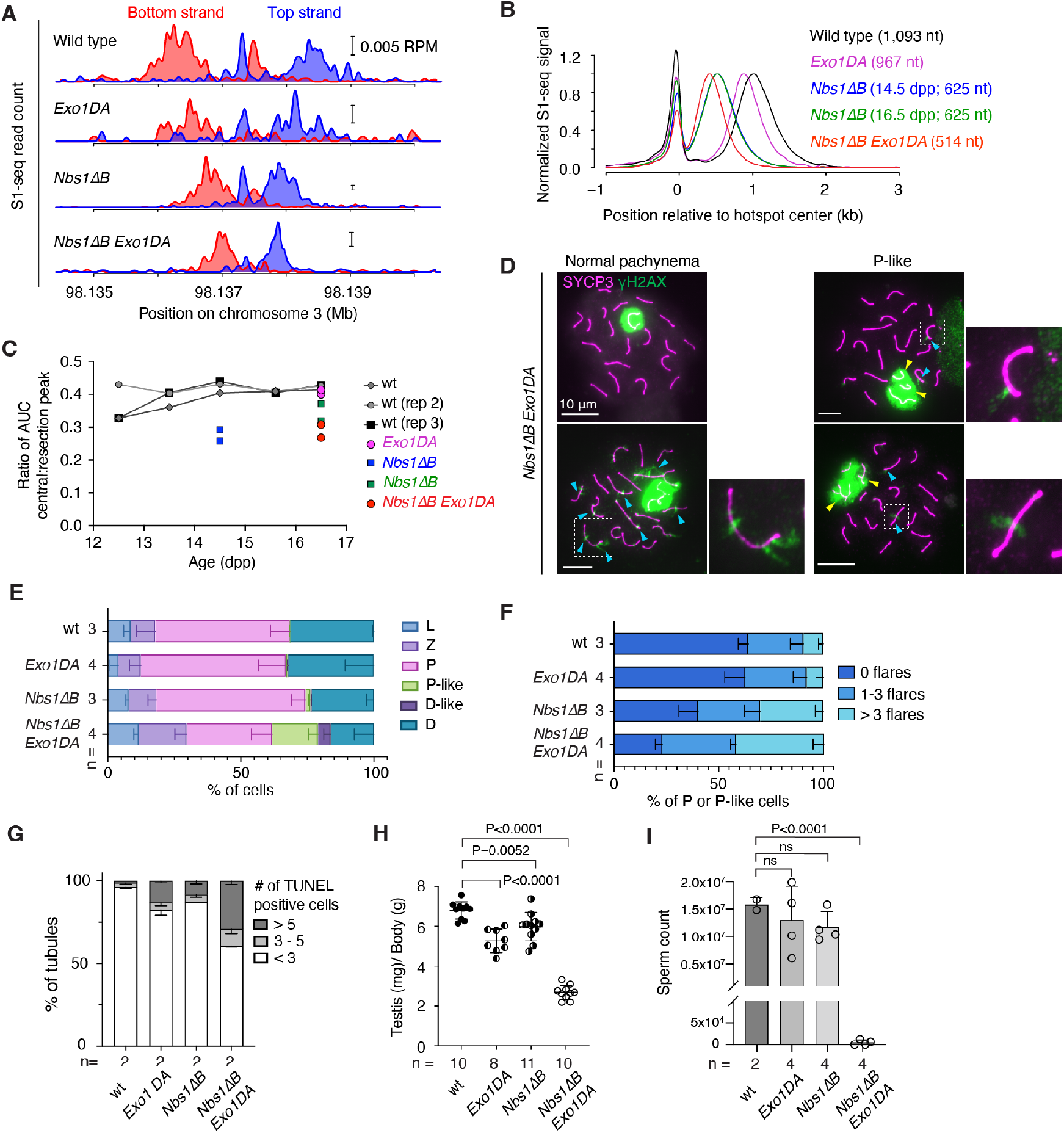
*Nbs1ΔB* and *Exo1DA* mutations affect resection additively. (A) S1-seq at the hotspot from **Figure 1F**. Because of the smaller size of *Exo1DA Nbs1ΔB* pups, we collected samples at 16.5 dpp for this experiment. Wild type (14.5 dpp) is reproduced from **Figure 1F**; two biological replicates were averaged for each of the other genotypes. (B) Genome-average S1-seq patterns at hotspots, smoothed with a 151-bp Hann window. Wild type and *Nbs1ΔB* (14.5dpp) are reproduced from **Figure 5D**. Mean resection lengths are indicated. (C) Reduced central S1-seq signal (presumed recombination intermediates) in *Nbs1ΔB* mutants. The plot shows the relative amount of central signal calculated as the area under the curve (AUC) for the central peak (–300 to +100 bp relative to hotspot centers) divided by the AUC for the resection peak (+100 to +2500 bp). For wild type, in each of three independent time courses, samples were collected from a single litter daily at approximately the same time each day. (D) Homolog pairing and/or synapsis defects. Representative spreads of *Nbs1ΔB Exo1DA* spermatocytes are shown with either normal-appearing SYCP3 and γH2AX staining or with unsynapsed or tangled chromosomes within the γH2AX-stained domain (yellow arrowheads). Blue arrowheads point to examples of γH2AX flares on synapsed autosomes. (E) Spermatocyte stages based on SYCP3 and γH2AX staining. Error bars indicate mean ± SD of the indicated number of animals. (F) Percentage of pachytene or pachytene-like cells with no, 1–3, or >3 γH2AX flares. Error bars indicate mean ± SD of the indicated number of animals. (G) Frequencies of tubules with the indicated number of TUNEL-positive cells. Error bars indicate mean ± range for the indicated number of animals. (H) Reduced testis size in *Nbs1ΔB Exo1DA* double mutants (5–11 wk old). Error bars indicate mean ± SD. P values are from Student’s t tests. (I) Sperm counts (mean ± SD). Results of Student’s t tests are shown.

Relative to the S1-seq signal from resection endpoints, the central signal from recombination intermediates appeared lower in *Nbs1ΔB Exo1DA* than in wild type or the single mutants (**Figure 7B**). To evaluate this, we measured total read count (area under the curve) separately for the central peak and for the resection endpoint peak and calculated the ratio between these values (**Figure 7C**). In wild type, the central peak was reproducibly ~40% of the level of the resected DSBs from 13.5 to 16.5 dpp. This ratio was lower at 12.5 dpp (~33%) in two of three experiments (**Figure 7C**). Since this earlier age is likely to be more enriched for earlier prophase I spermatocytes, the lower relative amount of central signal may reflect timing differences between DSB formation (appearance of the resected DSB signal) and formation of the recombination intermediates detected by S1-seq.

In *Nbs1ΔB* mice, the relative amount of central signal was substantially lower at 14.5 dpp (27.5% of the DSB signal; mean of n = 2) and it increased but remained lower than wild type at 16.5 dpp (34.7%; **Figure 7C**). The central signal at 16.5 dpp was even lower in *Nbs1ΔB Exo1DA* (28.8%) but the *Exo1DA* mutation by itself had no effect. These decreased yields of the central signal in *Nbs1ΔB* single or double mutants may indicate that fewer of the relevant recombination intermediates are formed, that they turn over more rapidly, and/or that they exhibit structural features that make them more difficult to detect by S1-seq. Regardless of the source of the changes, however, it appears that *Nbs1ΔB* mutants experience alterations in interhomolog recombination that are exacerbated when combined with *Exo1DA*.

Reinforcing this view, *Nbs1ΔB Exo1DA* double mutants displayed an elevated frequency of pachytene-like cells with synaptic failures (tangled and/or unpaired chromosomes) and a higher proportion of zygotene cells than normal, possibly indicative of delays in completing synapsis (**Figures 7D,E**). Pachytene and pachytene-like cells from the double mutant also showed an increased frequency of γH2AX flares on synapsed chromosomes, indicative of delays in DSB repair (**Figures 7D,F**). Seminiferous tubule sections showed few if any elongating spermatids in *Nbs1ΔB Exo1DA* double mutants (**Figure S7D**) and an elevated frequency of apoptotic spermatocytes (**Figures 7G and S7E**). As a consequence of this germ cell loss, the mice had substantially reduced testis sizes (**Figure 7H**) and were essentially devoid of epididymal sperm (**Figures 7I and S7D**). Single *Nbs1ΔB* or *Exo1DA* mutants showed few of these defects, consistent with previous studies^59,74^, although apoptotic tubules were observed at modestly elevated frequencies and testis sizes were reduced (**Figures 7G,H**).

## DISCUSSION

Unlike the paradigmatic budding yeast pathway for meiotic DSB resection^11,12^, mice rely less on EXO1^14,15^ and we now show that multiple MRN mutations confer drastic resection defects (see also^20^). Thus, while the canonical two-step handoff from MRN(X) to EXO1 occurs in both yeast and mammals, we find that mouse MRN is essential not just for resection initiation, but also unexpectedly for most long-range resection as well. Since *Exo1DA* combined additively with MRN deficiency, EXO1 appears to “polish” resection tracts that are generated primarily through MRN.

ATM deficiency decreases resection lengths for a subset of DSBs^14,15^. Thus, because the MRN hypomorphs compromise ATM activation^21,26,75^, their resection defects could be an indirect consequence of diminished ATM signaling. However, *Mre11-cHN* mice appear to have normal ATM activity^29,53,56,57^ (**Figure 4F**), suggesting that their defective resection is tied more directly to attenuated MRN activity per se. By extension, the resection defects in MRN hypomorphic adults and *Mre11-cKO* juveniles may be explained in a similar manner. We cannot exclude that reduced ATM activity also contributes, but *Mre11-ATLD1* and *Nbs1ΔB* mutants only modestly attenuate meiotic ATM activation as judged by their retention of most ATM-dependent DSB control^76^, so this explanation would require that resection is particularly sensitive to small decreases in ATM activation. Moreover, the resection-promoting function of ATM itself remains unknown, so the opposite causality is also plausible: i.e., resection defects in *Atm* mutants might hypothetically be an indirect consequence of losing ATM-dependent regulation of MRN.

We consider two plausible ways in which MRN might promote long-range resection aside from ATM activation. First, it might directly resect DNA via iterative nicking that spreads progressively from the DSB end. Although yeast MRX and human MRN preferentially cleave the 5’-terminal DNA strand ~15–45 nt away from a blocked DNA end in vitro^77-80^, MRN can both oligomerize and diffuse on DNA^81,82^. Moreover, MRX can resect unrepairable DSBs in vivo more than one kilobase in yeast lacking long-range resection enzymes^83^, and iterative nicking was proposed to explain both short-range resection in vegetative yeast cells^84^ and the ~300 nt average resection in meiotic cells lacking Exo1 activity^11^.

A second, non-exclusive possibility is that MRN recruits or activates another nuclease(s). MRN enhances DNA2 nuclease in vitro^68,85-88^, although it remains unclear if the same is true in vivo. CtIP also promotes DNA2 activity both in vitro and in vivo^89,90^. Budding yeast Dna2 is unlikely to be important for meiotic resection because its partner helicase Sgs1 is dispensable^10-12^. However, the possibly mammal-specific interaction between CtIP and DNA2^89,90^ may make DNA2 a candidate for long-range resection in mouse meiosis^15^.

In contrast to its unanticipated requirement in long-range resection, MRN involvement in resection initiation was expected. However, the specific genetic dependencies were intriguing. For example, we were surprised to observe only minor defects in resection initiation in *Mre11-cHN* mice. It is formally possible that MRE11 nuclease is dispensable for clipping SPO11 from DSBs even though MRE11 protein itself is required. However, we consider it more likely that this result instead reflects continuing presence of sufficient wild-type MRE11. For example, dimerization with continuously expressed MRE11-H129N might slow the degradation of wild-type protein compared to what happens in *Mre11-cKO*. In this scenario, mutant/wild-type heterodimers are still active for strand cleavage since they retain a functional nuclease active site. Alternatively, MRE11-H129N might amplify activity of a small amount of residual wild-type protein in a structural role via heterodimer formation or oligomerization on DNA. It is also possible that binding of inactive MRN complexes to DSBs allows another nuclease(s) to be recruited to initiate resection in an unconventional manner.

The *Mre11-cKO* mutant is the first example in mammals of a yeast *rad50S*-like meiotic phenotype, namely, accumulation of completely unresected DSBs^6,91-93^. This finding definitively establishes that MRN is essential for resection initiation. The elevated DSB sequencing signal and double cutting in this mutant also confirms and extends the previous conclusion, based on modest elevation of SPO11-oligo complexes in *Mre11-ATLD1 and Nbs1ΔB*, that MRN promotes ATM activation in the context of controlling SPO11 activity^76^. While this manuscript was in preparation, conditional deletion of *Rad50* (with *Stra8-Cre*) was also found to cause an increased frequency of unresected DSBs, but to a much more modest extent than we observed with *Mre11-cKO* and without an apparent increase in DSB levels or evidence of double-cutting^20^. Because resection was not directly assessed for *Nbs1* conditional deletion (with *Vasa-Cre*), it is unclear whether NBS1 is required for resection initiation^19^. The differences between the conditional mutants may reflect differences in timing and efficiency of gene deletion and protein disappearance rather than different requirements for the individual MRN subunits.

Our findings may also indicate that mouse MRE11 is dispensable for DSB formation, unlike in *S. cerevisiae* and *Caenorhabditis elegans*, but similar to *Schizosaccharomyces pombe, Drosophila melanogaster*, and *Arabidopsis thaliana* [reviewed in^94^]. However, this interpretation should be viewed cautiously because *Mre11-cKO* spermatocytes may retain trace amounts of MRE11 protein that are sufficient for DSB formation but not resection. In this vein, we note that *Mre11* and *Ctip* null mutations are both cell-lethal, whereas conditional deletion of *Mre11* but not of *Ctip* was tolerated by spermatogonia and premeiotic spermatocytes. This is consistent with both proteins being essential for mitotic germ cell divisions, but MRE11 protein persisting longer than CtIP after Ngn3-Cre-mediated excision.

In addition to defining the genetic control of DSB resection, our findings offer insight into the minimal resection length needed for successful downstream steps. The extensive pairing and synapsis defects in *Mre11-cKO* mice met the expectation that recombination cannot proceed without ssDNA generated by resection. However, our findings showed that meiosis is remarkably robust as long as some resection occurs. Recombination, chromosome pairing, synapsis, and chromosome segregation all occurred efficiently with slightly more than half the normal average resection length (*Mre11-ATLD1*). Severe problems were seen only in the *Nbs1ΔB Exo1DA* double mutant, with its only modest further decrement in resection length. But even in this mutant, although most cells failed to complete meiosis, most chromosomes paired and synapsed well, indicating that most DSBs were successfully engaging in recombination.

It is unclear why recombination begins to fail in the *Nbs1ΔB Exo-1DA* double mutant but not appreciably in the *Nbs1ΔB* or *Mre11-ATLD1* single mutants. The minimal length of ssDNA needed to support homology searching and strand exchange in mammalian meiosis is unknown, but vegetative yeast cells can execute recombination with less than 40 bp of homology, and normal recombination efficiency appears to need only~100–250 bp^95-99^. Moreover, RAD51 and DMC1 filaments are thought to occupy only a few hundred nucleotides^66,67^. Although the distribution of resection tract lengths is highly stereotyped across all mouse hotspots, there is substantial variation from DSB to DSB, and perhaps between the two sides of a single DSB. It has been suggested for yeast that normal resection lengths ensure that all DSBs are resected enough to support efficient, accurate recombination^12^. It is therefore possible that *Nbs1ΔB Exo1DA* double mutants become vulnerable to recombination defects because a large enough subset of their DSBs fall below the minimum length needed for strand exchange per se or for proper choice of homologous recombination partner^100^.

## MATERIALS AND METHODS

### Mice

Experiments conformed to the US Office of Laboratory Animal Welfare regulatory standards and were approved by the Memorial Sloan Kettering Cancer Center Institutional Animal Care and Use Committee. Mice were maintained on regular rodent chow with continuous access to food and water until euthanasia by CO_2_ asphyxiation prior to tissue harvest. Previously described *Mre11-flox* and *Mre11-H129N*^29^ and *Ctip-flox*^101,102^ alleles were crossed with *Ngn3-Cre*^30^ to create male germ-line-specific mutations.

To generate *Mre11-cKO* experimental mice, *Mre11*^*flox/flox*^ (or *Mre11*^*wt/flox*^ based on availability) mice were crossed with *Mre11*^*+/–*^ *Ngn3-Cre*^*+*^ mice. The *Mre11-deletion* allele (*Mre11*^*–*^) was derived by Cre-mediated recombination of the *Mre11-flox* allele. To generate *Mre11-cHN* experimental mice, *Mre11*^*H129N/+*^ *Ngn3-Cre*^*+*^ mice were crossed with *Mre11*^*flox/flox*^ mice. To generate *Ctip-cKO* mice, *Ctip*^*flox/flox*^ mice were crossed with *Ctip*^*+/–*^ *Ngn3-Cre*^*+*^ mice. The *Ctip* deletion allele (*Ctip*^*–*^) was derived by Cre-mediated recombination of *Ctip-flox*.

Although *Ngn3-Cre* is also expressed in embryonic pancreas and endocrine cells^30^, we did not observe any gross defects of mutant animals created with *Ngn3-Cre* up through the ages of testis sample collection. Previously described *Atm* (Barlow et al. 1996), *Spo11*^103^, *Rad50S*^23^, *Mre11-ATLD1*^21^, *Nbs1ΔB*^26^, and *Exo1DA* (Zhao et al. 2018) mutations were maintained on a congenic C57BL/6J strain background. *Nbs1ΔB Exo1DA* animals were generated by crossing *Nbs1ΔB* mice with *Exo1DA* mice. To enhance the survival rate of double homozygous male mice for experimental purposes, female pups were euthanized by hypothermia between 4–6-dpp in accordance with the approved animal protocol.

To generate targeted *Rad50* mutations, a guide RNA cassette with sequence (5’-TACATTTGTTCATGATCCCA(AGG)) and single-stranded donor DNA (5’TTATAAAAATTTAATTCTTAAAATA-CAAAACTTTACAGACACCATTACCTTGGGATaATGAACAAAT-GTATTTCCTTTGGTTCCAGGAGGGAAATCTCCAGTACAA)harboring desired missense mutations for D69>Y, or a guide RNA cassette with sequence (5’-TCTCGGTCGAGATTTGTTGT(CGG)) and single-stranded donor DNA (5’-GGAAACCTTCTGTCTGAACT-GTGGCATCCTTGCCTTGGATGAGCCGACAACAAATtttGAC-CGAGAAAACATTGAGTCTCTTGCACATGCTTTGGTTGAG-TAAGTA) harboring desired missense mutations for L1237>F were microinjected into pronuclei of zygotes. Founder mice were crossed to C57BL/6J mice purchased from Jackson Laboratories to obtain germline transmission, then heterozygous animals were backcrossed to C57BL/6J for at least three generations.

### S1-seq and Exo7/T-seq

#### Choice of sequencing method and animal ages

S1-seq was originally applied to juvenile testis samples because these samples lack post-meiotic cells and are thus enriched for spermatocytes that contain DSBs and recombination intermediates^15^. We continued to use juvenile samples for most experiments here, but found that Cre-mediated excision of the *Mre11-flox* construct was more penetrant in adult samples. Initial attempts to perform S1-seq on adult samples gave poor signal strength from SPO11-generated DSBs at hotspots because of the presence of S1-sensitive triplex secondary structures in the DNA^104^. We therefore initially used Exo7/T-seq for analysis of resection in *Mre11-cKO* and *Mre11-cHN* mice, because exonuclease VII and exonuclease T require DNA ends and thus do not give sequencing reads from many of the non-DSB structures that S1-seq can pick up. We later found that increasing the pH during the nuclease S1 digestion step greatly improved the specificity for SPO11-initiated events by avoiding the formation of triplex structures^34^, so we repeated the analysis of *Mre11-cKO* using S1-seq on adult samples. For the wild-type time courses (**Figure 7C**), each time point was processed up to the agarose plug step, then S1-seq maps were prepared in a single batch after collecting the last time point.

#### Library preparation and sequencing

Libraries were prepared as described elsewhere^34^, based on S1-seq^15^ and END-seq^14,105^. Briefly, testes from mice at the indicated ages were decapsulated and digested with collagenase type IV (Worthington) and Dispase II (Sigma). The resulting seminiferous tubule preparations were then further treated with TrypLE Express enzyme (GIBCO) and DNase I (Sigma) for 15 min at 35 °C. TrypLE enzyme was inactivated with 5% FBS and tubules were further dissociated by repeated pipetting. Cells were then passed through a 70-μm cell strainer (BD Falcon).

One to two million cells from juvenile testes or two to three million cells from adult testes were embedded in plugs of 1% low-melting-point agarose (Lonza). After a brief incubation at 4 °C until the agarose became solid, plugs were incubated with proteinase K (Roche) in lysis buffer (0.5 M EDTA at pH 8.0, 1% N-lauroylsarcosine sodium salt) at 50 °C over two nights. Plugs were washed five times for 20 min with TE (10 mM Tris-HCl at pH 7.5, 1 mM EDTA at pH 8.0), and then incubated with 100 μg/ml RNase A (Thermo) for 3 hr at 37 °C. Plugs were then washed five times with TE and stored in TE at 4 °C until usage.

All library types were prepared in the same way aside from the ssDNA digestion step. For S1-seq, plugs were equilibrated with S1 buffer (50 mM sodium acetate, 280 mM NaCl, 4.5 mM ZnSO_4_, pH 4.7 at 25 °C),then treated with 9 U of nuclease S1 (Promega) for 20 min at 37 °C. For Exo7/T-seq, plugs were instead equilibrated with exonuclease VII buffer (50 mM Tris-HCl, 50 mM sodium phosphate, 8 mM EDTA, 10 mM 2-mercaptoethanol, pH 8 at 25 °C), then treated with 50 U of exonuclease VII (NEB) for 60 min at 37 °C. Plugs were then equilibrated in NEBuffer 4 and treated with 75 U of exonuclease T (NEB) for 90 min at 24 °C.

Plugs were next equilibrated with T4 polymerase buffer (T4 ligase buffer (NEB) supplemented with 100 μg/ml BSA and 100 μM dNTPs (Roche)) then incubated with 30 U T4 DNA polymerase (NEB) at 12°C for 30 min. Plugs were then washed in TE and equilibrated in T4 ligase buffer on ice. Biotinylated P5 adaptors were ligated to the blunted ends with 2000 U T4 DNA Ligase (NEB) at 16 °C for 20 hr. After ligation, plugs were soaked in TE overnight at 4 °C to diffuse excess unligated adaptors out of the plugs. The plugs were then equilibrated in 1× β-agarase I buffer (10 mM Bis-Tris-HCl, 1 mM EDTA, pH 6.5), incubated at 70 °C for 15 min to melt the agarose with brief vortexing every 5 min, mixed well, cooled to 42 °C, and digested with 2 μl of β-agarase I (NEB) at 42 °C for 90 min. DNA was sheared to fragment sizes ranging 200–500 bp with a Covaris system (E220 Focused-ultrasonicator, microtube-500) using the following parameters: delay 300 s then three cycles of [peak power 175 W, duty factor 20%, cycles/burst 200, duration 30 s, and delay 90 s]. The DNA was precipitated with ethanol, then dissolved in TE. Unligated adaptors were removed with SPRIselect beads (Beckman Coulter). Fragments containing the biotinylated adaptor were further purified with Dynabeads M-280 streptavidin (Thermo) and DNA ends were repaired using the End-it DNA End-repair kit (Lucigen). P7 adaptors were ligated to DNA fragments, then PCR was done directly on the bead-immobilized fragments. PCR products were purified with AMPure XP beads (Beckman Coulter) to remove primer dimers and unligated adaptors.

After Qubit (Thermo) quantification and quality control by Agilent BioAnalyzer, libraries were pooled and sequenced on the Illumina NextSeq, HiSeq, or NovaSeq platforms in the Integrated Genomics Operation (IGO) at Memorial Sloan Kettering Cancer Center. A spike-in of bacteriophage FX174 DNA was added to increase diversity when necessary and for quality control purposes. We obtained paired-end reads of 50 bp.

#### Preprocessing and mapping

Base calls were performed using bcl-convert software v.3.9.3 to 3.10.5 or bcl2fastq software v.2.20. Reads were trimmed and filtered by Trim Galore version 0.6.6 with the arguments --paired --length 15 (http://www.bioinformatics.babraham.ac.uk/projects/trim_galore/). Sequence reads were mapped onto the mouse reference genome (mm10) by bowtie2 version 2.3.5.1^106^ with the arguments-N 1 -X 1000. Duplicated reads were removed by Picard (https://broadin-stitute.github.io/picard/). Uniquely and properly mapped reads (MAPQ ≥ 20) were extracted by samtools with the argument -q 20 (http://www.ht-slib.org/). Reads were assigned to the nucleotide immediately next to the biotinylated adaptor. Mapping statistics are in **Supplemental Table S2**.

#### Bioinformatic analysis and plotting

Maps were analyzed using R version 4.0.3 (http://www.r-project.org). To generate genome-average profiles around hotspots (e.g., **Figure 4D**), an estimated background was removed by subtracting from all values the value of signal 2,500 bp away from the hotspot center. The signal was then normalized to the peak height of resection endpoints, i.e, the maximum value among positions from 100 to 2,500 bp. Negative values were set as zero for plotting purposes. Where appropriate, data were smoothed with a Hann function. To calculate the mean resection length, S1-seq signal was averaged across hotspots and an estimated background was removed by subtracting from all values the value of signal 2.5 kb away from the hot spot center. The signal close to and further away from the hot spot center was excluded by setting values of positions <100 bp and >2.5 kb to zero. Fractions of total signal were calculated every 100 bp and the mean resection length was calculated. The modal resection length in **Figure S2F** was determined by the peak position of the genome-average profile between 100 bp and 2,500 bp from hotspot centers.

The color scale of heatmaps were defined after local normalization of sequencing signal at each hotspot. Signals in 40-bp bins were divided by the total signal in a 4,001-bp window around each hotspot center, so that each row has a total value of 1 regardless of the strengths of hotspots and color reflects the local spatial pattern. Normalized signals were classified into 10 groups by deciles and color-coded.

### MRE11 ChIP-seq

MRE11 ChIP was performed following the previously published DISCOVER-seq protocol^107^. In brief, fresh testes of 5–7 wk old wild-type, *Atm*^*–/–*^, or *Mre11-cHN* mice were dissociated into single cells and resuspended in DMEM. PFA (16% stock, Pierce) was added to a final concentration of 1% and incubated for 15 min with gentle agitation, then the crosslinking was stopped by adding 2.5 M glycine to a final concentration of 125 mM, followed by 3 min incubation on ice. The nuclei were prepared as described^107^ and up to 10 million nuclei were resuspended in 500 μl lysis buffer (0.5% *N*-lauroylsarcosine, 0.1% sodium deoxycholate,10 mM Tris-HCl pH 8.0, 100 mM NaCl, 1 mM EDTA) supplemented with protease inhibitor (Roche), and sonicated to fragment sizes ranging 200–300 bp with a Covaris system (E220 Focused-ultrasonicator, microtube-500) using the following parameters: delay 300 s then six cycles of [peak power 75 W, duty factor 10%, cycles/burst 200, duration 150 s, and delay 60 s]. Sheared nuclei were centrifuged at 20,000 *g* for 10 min at 4°C to remove debris and Triton-X (1% (vol/vol) final concentration) and lysis buffer were added to make the total volume 3 ml.

For each immunoprecipitation, 10 μl of anti-MRE11 (Novus Biologicals NB100-142) was pre-mixed with 100 μl protein A Dynabeads, then added to chromatin lysates and incubated overnight at 4 °C with end-over-end rotation. Immunoprecipitated DNA was eluted and crosslinks were reversed as described^107^. ChIP sequencing libraries were prepared from cleaned DNA as described for SSDS libraries^108^ without the boiling step after the end-repair and dA-tailing steps. After ligation with sequencing adaptors, the library was amplified by 15 cycles of PCR. The amplified library was purified and sequenced on the Illumina NovaSeq platform in the IGO. We obtained paired-end reads of 100 bp.

Base calls were performed using bcl-convert software v.3.9.3 to 3.10.5 or bcl2fastq software v.2.20. Sequencing reads were processed and mapped as described for resection libraries and uniquely and properly mapped reads were counted, and maps were further analyzed using R version 4.0.3. Mapping statistics are in **Supplemental Table S2**.

### Single-stranded DNA sequencing (SSDS) library preparation and data analysis

ChIP followed by SSDS was performed as previously described^64,108^. In brief, a decapsulated, fresh or fresh-frozen testis of 8-12 wk old wild-type or *Nbs1ΔB* mice was placed in room temperature 1% PFA (16% stock, Pierce) in PBS and incubated for 15 min with gentle agitation. The crosslinking was stopped by adding 2.5 M glycine to a final concentration of 125 mM, followed by 5 min incubation at room temperature. Fixed cells were immediately homogenized in a Dounce homogenizer with 20 strokes of the tight-fitting pestle, then passed through a 70 μm cell strainer. The nuclei were prepared as described^108^, resuspended in 900 μl shearing buffer (0.1% SDS, 10 mM Tris-HCl pH 8.0, 1 mM EDTA) supplemented with protease inhibitor (Roche), split into two~500 μl aliquots and sonicated to fragment sizes ranging 500–1000 bp with a Covaris system (E220 Focused-ultrasonicator, microtube-500) using the following parameters: delay 300 s then three cycles of [peak power 75 W, duty factor 10%, cycles/burst 200, duration 150 s, and delay 60 s]. Sheared nuclei were centrifuged at 12,000 *g* for 10 min at 4 °C to remove debris. The sonicated chromatin was then transferred to a 3 ml Slide-A-Lyzer G2 dialysis cassette (Pierce) and dialyzed with ChIP buffer (16.7 mM Tris-HCl, pH 8.0, 167 mM NaCl, 1.2 mM EDTA, 0.01% SDS, 1.1% Triton X-100) for 5 hr at 4 °C with constant but slow stirring. For each immunoprecipitation, 20 μl of anti-DMC1 (Abcam, ab11054), 10 μl of anti-RAD51 (Novus Biologicals, NB100-148), or 10 μl of anti-RPA2 (Abcam, ab76420) was added to chromatin lysates and incubated overnight at 4 °C with end-over-end rotation.

As described previously^108^, protein G or A Dynabeads were added to the chromatin and antibody mixture and incubated 2 hr at 4 °C, then immunoprecipitated DNA was washed and eluted. To reverse DNA-protein crosslinks, 12 μl of 5 M NaCl was added to 300 μl of eluted sample and incubated overnight at 65 °C. DNA was cleaned with MinElute PCR Cleanup Kit (Qiagen) after proteinase K (Thermo) treatment. Following end repair and dA-tailing, DNA was incubated at 95 °C for 3 min and cooled to room temperature to enrich for single-stranded DNA. After ligation with sequencing adaptors, SSDS library was amplified with 12 cycles of PCR. The amplified library was purified and sequenced on the Illumina NovaSeq platform in the IGO. We obtained paired-end reads of 100 bp.

Base calls were performed using bcl-convert software v.3.9.3 to 3.10.5 or bcl2fastq software v.2.20. We then ran a previously described bioinformatic pipeline on the resulting reads for identification of single-stranded sequences^108^. Sequence reads at SPO11-dependent hotspots were analyzed using R version 4.0.3.

### Sperm counts

Sperm isolation was performed as described^109^. Briefly, both caudal and caput epididymis were dissected from >5 wk old mice, with all excess fat and tissue trimmed away, and placed on top of filter mesh of 5 ml tubes with cell-strainer caps (BD) filled completely with PBS. After 5 min, the cap was carefully lifted out of the PBS to break the fluid surface tension, releasing plumes of sperm into the tube. This was repeated until sperm were no longer released after lifting out the cap. The cap (and tissue) was removed, and the tube sealed with parafilm. The sperm were pelleted in a swinging bucket rotor for 2 min at 4,000 *g*. The supernatant was aspirated down to ~1 ml, and then gently resuspended by quick vortex pulses and incubated for 1 min at 60 °C. Inactivated sperm were mixed with Trypan Bule solution and placed on a hematocytometer for counting.

### Spermatocyte chromosome spreads and immunostaining

Spreads were prepared as described with minor modifications^110^. When preparing a sequencing library and spreads from testes collected at the same time, testis cells were dissociated following the procedures in library preparation until passage through the 70 µm cell strainer (see above), then continued as described below. Briefly, decapsulated testis samples were placed in 2 ml TIM buffer (104 mM NaCl, 45 mM KCl,1.2 mM MgSO_4_, 0.6 mM KH_2_PO_4_, 0.1% glucose, 6 mM sodium lactate, 1 mM sodium pyruvate, pH 7.3), with 200 µl collagenase (20 mg/ml), then shaken at 350 rpm for 15 min at 32 °C. After washing, tubules were resuspended in 2 ml TIM buffer with 200 µl trypsin solution (7 mg/ml) and 20 µl DNase I (400 µg/ml) and shaken for 15 min at 350 rpm at 32 °C. To stop trypsin digestion, 500 µl of FBS was added. Resuspended cells were filtered through a 70 µm cell strainer and TIM buffer was added to 15 ml. After three washes with DNase I, cells were resuspended in 10 ml of TIM buffer and distributed in 1 ml aliquots to Eppendorf tubes, centrifuged, and the supernatant was removed. The cell pellet was resuspended in 80 µl pre-warmed 0.1 M sucrose and incubated for 10 min. From the cell suspension, 40 µl was added to a Superfrost glass slide covered with 130 µl 1% PFA (with 0.1% Triton X-100, pH 9.2) in a humid chamber. Slides were kept in the closed humid chamber for at least 2 hr at 4 °C, then air dried and rinsed with 0.4% Photo-Flo (Kodak). After air drying, slides were wrapped in aluminum foil and stored at −80 °C.

For immunofluorescence, slides were blocked for 10 min with dilution buffer (0.2% BSA, 0.2% fish skin gelatin, 0.05% Tween-20 in 1× PBS) at room temperature. Using a humid chamber, slides were incubated with 100 µl primary antibody solution overnight at 4 °C. After washing, secondary antibody was added for 1 hr at 37 °C. After washing, slides were mounted with Vectashield containing DAPI (H-1000, Vector Laboratories). Primary and secondary antibodies are listed in **Supplemental Table S3**.

### Histology

Testes and epididymides dissected from adult mice were fixed in Bouin’s fixative for 4 to 5 hr at room temperature, or in 4% PFA overnight at 4 °C. Bouin’s fixed testes were washed in 15 ml milli-Q water on a horizontal shaker for 1 hr at room temperature, followed by five 1-hr washes in 15 ml of 70% ethanol on a roller at 4 °C. PFA-fixed tissues were washed four times for 5 min in 15 ml milli-Q water at room temperature. Fixed tissues were stored in 70% ethanol before embedding in paraffin and sectioning at 5 μm. The tissue sections were deparaffinized with EZPrep buffer (Ventana Medical Systems). Antigen retrieval was performed with CC1 buffer (Ventana Medical Systems). Sections were blocked for 30 min with Background Buster solution (Innovex), and then avidin-biotin blocked for 8 min (Ventana Medical Systems). Hematoxylin and eosin (H&E) staining and immunohistochemical TUNEL assays were performed by the MSK Molecular Cytology Core Facility using the Autostainer XL (Leica Microsystems, Wetzlar, Germany) automated stainer for H&E or PAS with hematoxylin counterstain, and using the Discovery XT processor (Ventana Medical Systems, Oro Valley, Arizona) for TUNEL. Testis sections were incubated with anti-DDX4 or anti-MRE11 for 5 hr, followed by 60 min incubation with biotinylated goat anti-rabbit (Vector Labs) at 1:200 dilution (**Supplemental Table S3**). Detection was performed with DAB detection kit (Ventana Medical Systems) according to manufacturer’s instructions. Slides were counter-stained with hematoxylin and coverslips were mounted with Permount (Fisher Scientific).

### Image acquisition and analysis

Images of spread spermatocytes were acquired on a Zeiss Axio Observer Z1 Marianas Workstation, equipped with an ORCA-Flash 4.0 camera, illuminated by an X-Cite 120 PC-Q light source, with 100× 1.4 NA oil immersion objective. Marianas Slidebook (Intelligent Imaging Innovations, Denver Colorado) software was used for acquisition. Whole histology slides were scanned and digitized with the Panoramic Flash Slide Scanner (3DHistech, Budapest, Hungary) with a 20× 0.8 NA objective (Carl Zeiss, Jena, Germany). High-resolution histology images were acquired with a Zeiss Axio Imager microscope using a 63× 1.4 NA oil immersion objective (Carl Zeiss, Jena, Germany). All images were processed in Fiji^111^. To quantify foci of RPA, DMC1, or RAD51, staging of spermatocytes was assessed by SYCP3 staining (% of axes synapsed)and only foci colocalizing with chromosome axis (SYCP3-signal) were manually counted. Statistical analysis was done with Graphpad Prism 9.

### Structured illumination microscopy

Spermatocytes from juvenile *Nbs1ΔB* homozygous and heterozygous littermates (13–15 dpp) were prepared for surface spreading and subsequent immunofluorescence as described previously^64,112^. Briefly, testes were dissected and placed in 1× PBS, pH 7.4 at room temperature before removal of the tunica albuginea. Seminiferous tubules were incubated in hypotonic extraction buffer (30 mM Tris-HCl pH 8.2, 50 mM sucrose, 17 mM trisodium citrate dihydrate, 5 mM EDTA, 2.5 mM DTT, 1 mM PMSF) for 20 min at room temperature. A homogeneous cell suspension was made in 100 mM sucrose and fixed on slides covered with a 1% PFA solution (pH 9.2) containing 0.15% Triton X-100 for at least 3 hr in a humid chamber. Slides were either used for immunofluorescence staining immediately or stored at –80 °C. For immunofluorescence, slides were blocked for 30 min at room temperature in PBST (1x phosphate-buffered saline (PBS) containing 0.1% Triton X-100) containing 3% nonfat milk. Slides were incubated with primary antibodies overnight in a humid chamber at 37 °C. Slides were washed 3 × 10 min in PBST, then incubated with secondary antibody for 1 hr at 37 °C in a humid chamber. Both primary and secondary antibodies were diluted in PBST containing 3% nonfat milk. After secondary antibody incubation, slides were washed 3 × 10 min in the dark with PBST and the slides were mounted with VECTASHIELD mounting medium containing DAPI (H-1000, Vector Laboratories).

Structured illumination microscopy (3D-SIM) was performed at the Bio-Imaging Resource Center in Rockefeller University using an OMX Blaze 3D-SIM super-resolution microscope (Applied Precision), equipped with 405 nm, 488 nm and 568 nm lasers, and 100× 1.40 NA UPLSAPO oil objective (Olympus). Image stacks of several μm thickness were taken with 0.125 μm optical section spacing and were reconstructed in Deltavision softWoRx 7.0.0 software with a Wiener filter of 0.002 using wavelength specific experimentally determined OTF functions. Maximum intensity projection images were acquired in Deltavision softWoRx 7.0.0 software. Data presented in this study were pooled from two independent staining experiments. To ensure correct alignment of fluorescent channels in all SIM experiments, multi-color fluorescent beads (TetraSpeck microspheres, 100 nm, Life Technologies, T7279) were imaged and used to correct shifts from passage of different fluorescence wave lengths through the optical path, as previously described^64^. SIM image analyses were as previously described^64^.

### Immunoblotting

About one million dissociated testis cells were boiled with NuPAGE LDS Sample Buffer (Life Technologies) and separated on 3%–8% Tris-acetate NuPAGE precast gels (Life Technologies) at 150 V for 70 min. Proteins were transferred to polyvinylidene difluoride (PVDF) membranes by wet transfer method in Tris-glycine-20% methanol, at 120 V for 40 min at 4°C. Membranes were blocked with 5% non-fat milk in 1’ Tris buffered saline, 0.1% Tween (TBS-T) for 30 min at room temperature on an orbital shaker. Blocked membranes were incubated with primary antibodies overnight at 4°C. Membranes were washed with TBS-T for 30 min at room temperature, then incubated with HRP-conjugated secondary antibodies for 1 hr at room temperature. Membranes were washed with TBS-T for 30 min and the signal was developed by ECL Prime (GE Healthcare). Primary and secondary antibodies are listed in **Supplemental Table S3**.

### SPO11-oligo complexes and denaturing PAGE

We purified SPO11-oligo complexes from testes of adults (> 5 wk old) as previously described^110^. Briefly, testes were decapsulated, flash frozen in liquid nitrogen and stored at –80 °C. Testes were homogenized and extracts were cleared by ultracentrifugation. SPO11 was then immunoprecipitated with mouse monoclonal anti-SPO11-antibody 180 and protein A-agarose beads. SPO11 was eluted in SDS-containing buffer then subjected to a second round of immunoprecipitation with the same antibody. The material from the second immunoprecipitation was radio-labeled with terminal deoxynucleotidyl transferase and [α-^32^P]-CTP then eluted with Laemmli sample buffer and separated by SDS-PAGE. Proteins were transferred to PVDF membrane. Alternatively, the size distribution of SPO11 oligos purified from two adult *Atm-KO* or *Mre11-cKO* mice was determined by protease digestion followed by radiolabeling with [α-^32^P]-GTP followed by electrophoresis on a 15% denaturing polyacrylamide gel, which was then dried on filter paper as previously described^113^. Radio-labelled species were detected with Typhoon phosphor imager after 48 hr exposure with Fuji phosphor screens, then quantified in Fiji^111^.

### Cell culture

Mouse embryonic fibroblasts (MEFs) were generated and cultured as described^23^. Mitotic index was quantified using early passage primary MEFs by measuring Ser10 phosphorylation of histone H3 by flow cytometry 1 hr after 3 Gy of IR exposure. Cells were treated with 10 μM of ATM inhibitor (KU55933, 11855, Sigma) 30 min before IR exposure as an ATM deficient control. Two independent MEF cell lines for either wild type or Rad50-D69Y were tested. For the colony formation assay, cells were immortalized as described^23^, then plated with different doses of CPT for 24 hr and washed and cultured in drug-free media. For ATR inhibition, cells were treated with 50 nM of ATR inhibitor (VE822, S7102, Selleckchem) 2 hr before CPT treatment. After 10 days of growth, colonies were visualized by crystal violet stain (0.5% crystal violet, 25% methanol).

## Data Availability

Raw and processed sequencing S1-, Exo7/T-seq, MRE11-ChIP, and SSDS data have been deposited in the Gene Expression Omnibus (GEO) repository under accession numbers GSE266258, GSE266151, and GSE266271. We used additional S1-seq and Exo7/T-seq data from GEO accession numbers GSE265863 and GSE229450^34,114^. We used SPO11-oligo data from GEO accession numbers GSE84689 and *Atm*^*–/–*^ MRE11 ChIP-seq data from GEO accession GSE138915^14,37^.

## ACKNOWLEDGMENTS

This article is subject to the Open Access to Publications policy of the Howard Hughes Medical Institute (HHMI). HHMI lab heads have previously granted a nonexclusive CC BY 4.0 license to the public and a sub-licensable license to HHMI in their research articles. Pursuant to those licenses, the author-accepted manuscript of this article can be made freely available under a CC BY 4.0 license immediately upon publication.

We thank W. Edelmann (Albert Einstein College of Medicine, New York, NY), R. Baer (Columbia University, New York, NY), Francesca Cole (MD Anderson Cancer Center, Houston, TX) and D. O. Ferguson (University of Michigan School of Medicine, Ann Arbor, MI) for generously sharing mouse lines. We thank A. Lukaszewicz (Univ. Michigan) and M. Jasin (MSK) for discussion. We thank the MSK Molecular Cytology core facility (N. Fan and M. Pulina) for histology; the MSK Integrated Genomics Operation (IGO) for sequencing; and the MSK Mouse Genetics Colony Management Group for assistance with mouse husbandry. MSK core facilities are supported by National Cancer Institute cancer center support grant P30 CA08748. The IGO was further funded by the Cycle for Survival and the Marie-Josée and Henry R. Kravis Center for Molecular Oncology. We thank M. Arter, E. Suranyi, H. Murakami, A. Shabro, V. Macera, and other members of the Keeney laboratory for discussions and experimental advice. This work was supported by NIH grants R35 GM118092 (to S.Keeney), R01 GM59413 (to J.H.J.P.), R35 GM136278 (to J.H.J.P.), and the Brain Pool Program through the National Research Foundation of Korea (NRF) funded by the Ministry of Science and ICT (2022H1D3A2A01096332 to S. Kim).

