## Supplemental material for "The MRE11-RAD50-NBS1 complex both starts and extends DNA end resection in mouse meiosis"

**This pdf contains:**

Supplementary Figures 1–7
Supplementary Tables 1–3

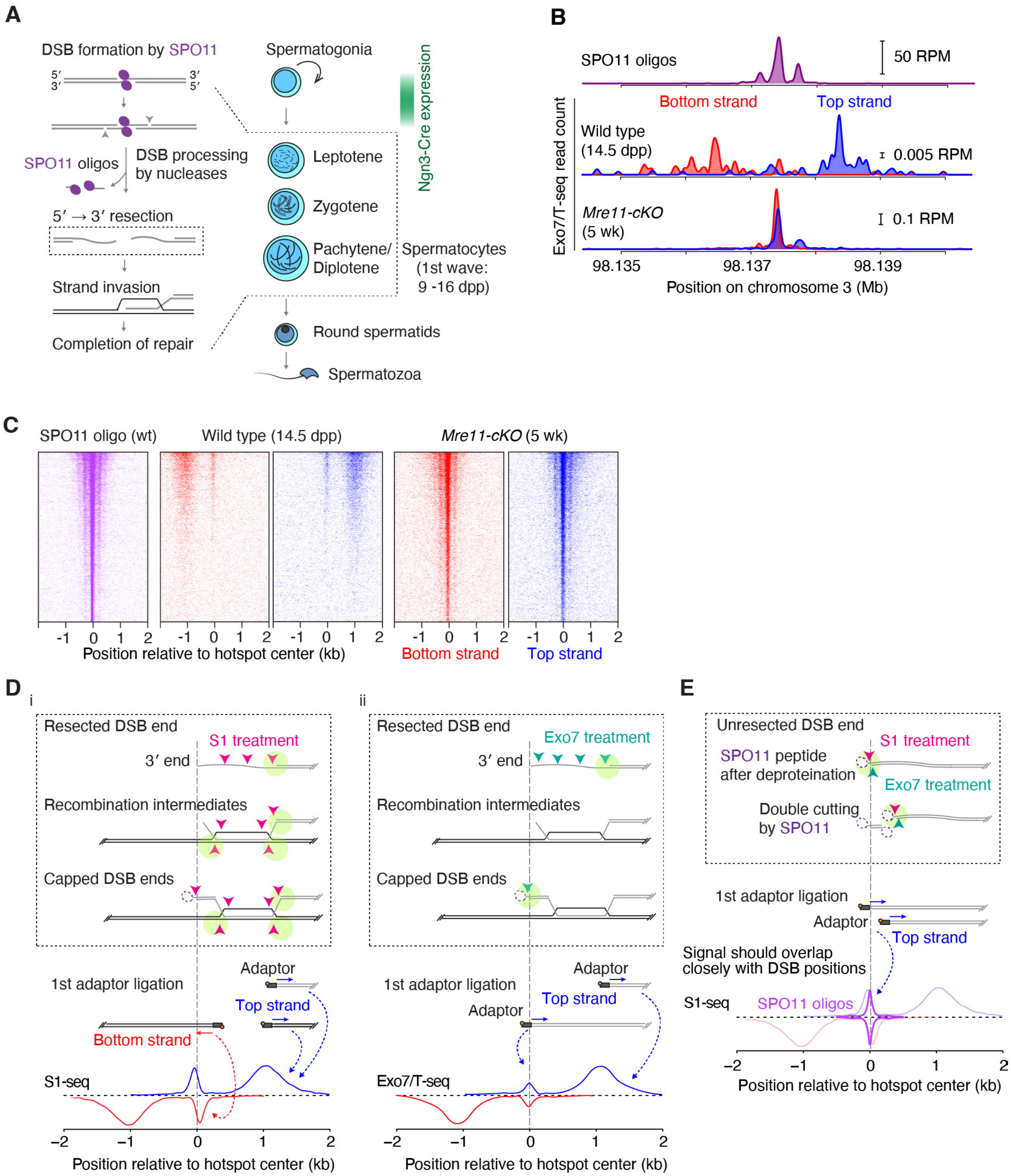


**Supplementary Figure 1. Resection defects in MRE11-depleted spermatocytes and comparison of S1-seq and Exo7/T-seq.**

(A) Schematic showing the timing of meiotic recombination steps (left) relative to the stages of spermatogenesis (right) and expected time of *Ngn3-Cre* expression.

(B,C) Confirmation using Exo7/T-seq that DSBs remain unresected in *Mre11-cKO* mice. Exo7/T-seq read distributions are shown for a representative hotspot (B) and around all hotspots (C). Heatmaps are presented as in **Figure 1G**.

(D) Schematics illustrating presumed cleavage by nuclease S1 (i) or exonuclease VII (ii) of resected DSBs, recombination intermediates, or SPO11-oligo-capped DSBs. The vertical

dashed lines align the different elements of the cartoons by the 3′ end of the DSB ssDNA tail. SPO11 cuts DNA with a 2-nt 5′ overhang and stays covalently bound to the overhang until released from DSB ends by resection. The existence of a SPO11-bound (capped) recombination intermediate was proposed previously[14,15]. During library preparation, deproteination by proteinase K removes the DNA-bound SPO11 from capped ends, leaving a small peptide adduct (dashed magenta ellipses). (i) The endonuclease activity of S1 cleaves ssDNA from resected and capped breaks in addition to D-loop recombination intermediates (pink arrowheads) and generates duplex ends suitable for sequencing adaptor ligation (pale green circle). During later steps of library preparation, the short duplex oligonucleotides produced from capped ends (as well as from D-loops) are not recovered[15]. Therefore, S1-seq primarily generates reads from resection endpoints, which are distributed approximately 1 kb away from the center of hotspots with the expected polarity. In addition, S1-seq generates reads from recombination intermediates near the hotspot center (referred to here as the central signal) that have opposite polarity. (ii) Exonuclease VII is a ssDNA specific exonuclease that degrades overhangs in either the 3'-to-5' or 5'-to-3' direction from a DNA terminus (green arrowheads), producing 4–10 nt oligonucleotides[115,116]. Exonuclease VII can also remove peptide blocks such as SPO11 or aborted topoisomerase II from DNA ends and can generate blunt DNA ends when combined with exonuclease T in the END-seq procedure[14,117]. Neither exonuclease VII nor exonuclease T is expected to be able to digest D-loops or similar structures, so the central signals from Exo7/T-seq are thought to be instead from SPO11-oligo-capped structures. Therefore, Exo7/T-seq gives a weaker central signal that is more congruent with DSB positions defined by SPO11 oligos.
(E) Schematics illustrating presumed cleavage by nuclease S1 or exonuclease VII of unresected DSBs.
Panels D and E are adapted from[32] under a CC-BY license.

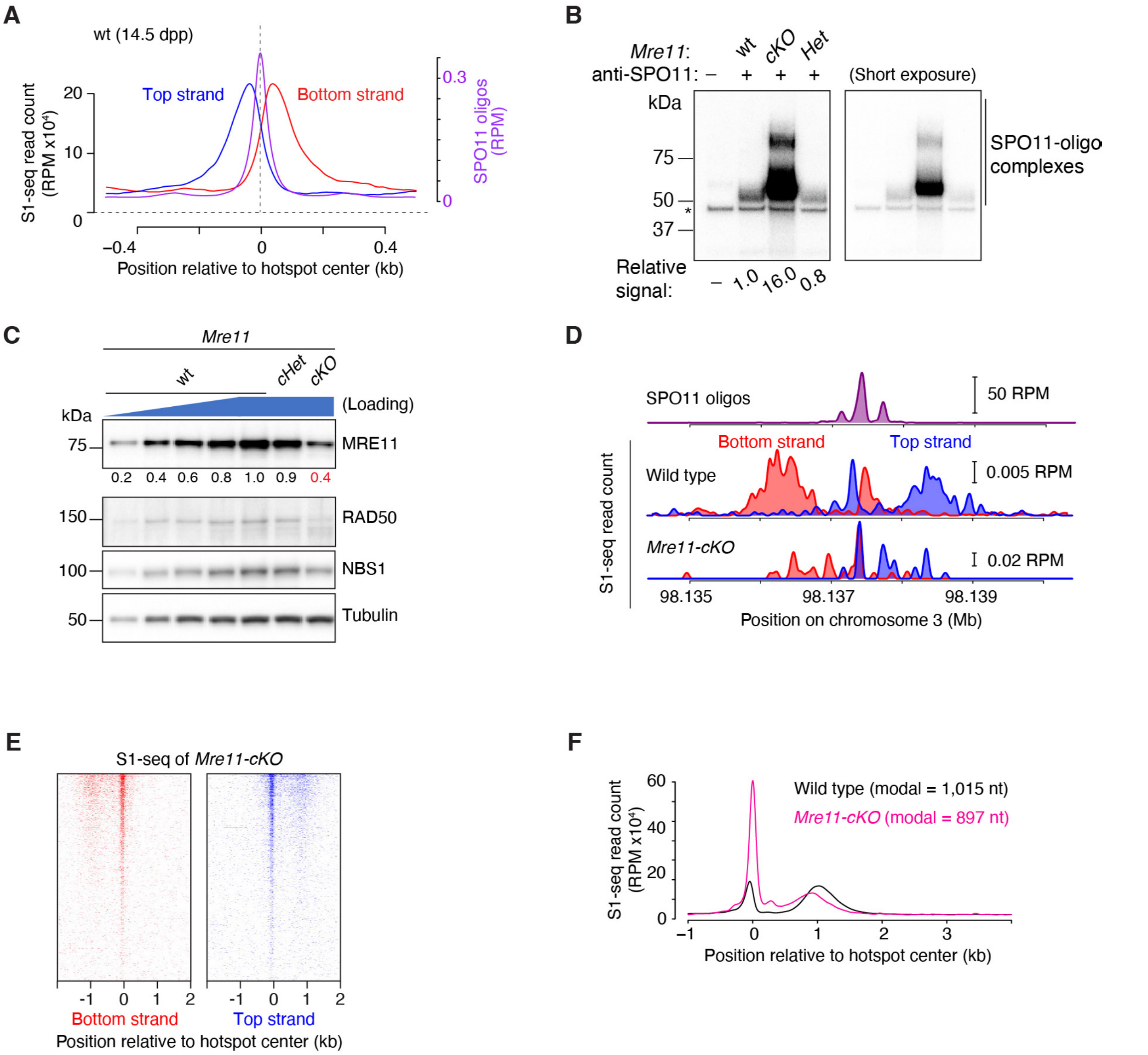


**Supplementary Figure 2. Further analysis of resection defects in MRE11-depleted spermatocytes**

(A) The central signal from recombination intermediates in wild type is distinct from the signal from unresected DSBs in *Mre11-cKO*. The genome-wide average of strand-specific S1-seq around hotspot centers in wild type is shown here, for comparison with the *Mre11-cKO* pattern in **Figure 2D**.

(B) Greatly elevated amounts of SPO11-oligo complexes in *Mre11-cKO*. A biological replicate is shown for the experiment in **Figure 2E**.

(C) MRN protein levels in juvenile *Mre11-cKO* mice. Immunoblots of whole-testis extracts from 14.5-dpp mice are shown as in **Figure 1C**.

(D) S1-seq signal at a representative DSB hotspot in 14.5-dpp *Mre11-cKO* mice (same hotspot as in **Figure 1F**). Wild type is reproduced from **Figure 1F** to facilitate comparison.

(E) Heatmaps of S1-seq signals around DSB hotpots from 14.5-dpp *Mre11-cKO* mice. Plots were generated as described in **Figure 1G**.

(F) Global average S1-seq profiles around hotspots from 14.5-dpp mice. Note the mix of unresected and resected DSBs in juvenile *Mre11-cKO* mice. We report modal resection lengths here instead of the mean values reported elsewhere because the high signal from unresected

DSBs in *Mre11-cKO* distorts the estimate of resection length. Wild type is reproduced from **Figure 2A**; three biological replicates for wild type and two for *Mre11-cKO* were averaged. Data are smoothed with a 151-bp Hann window.

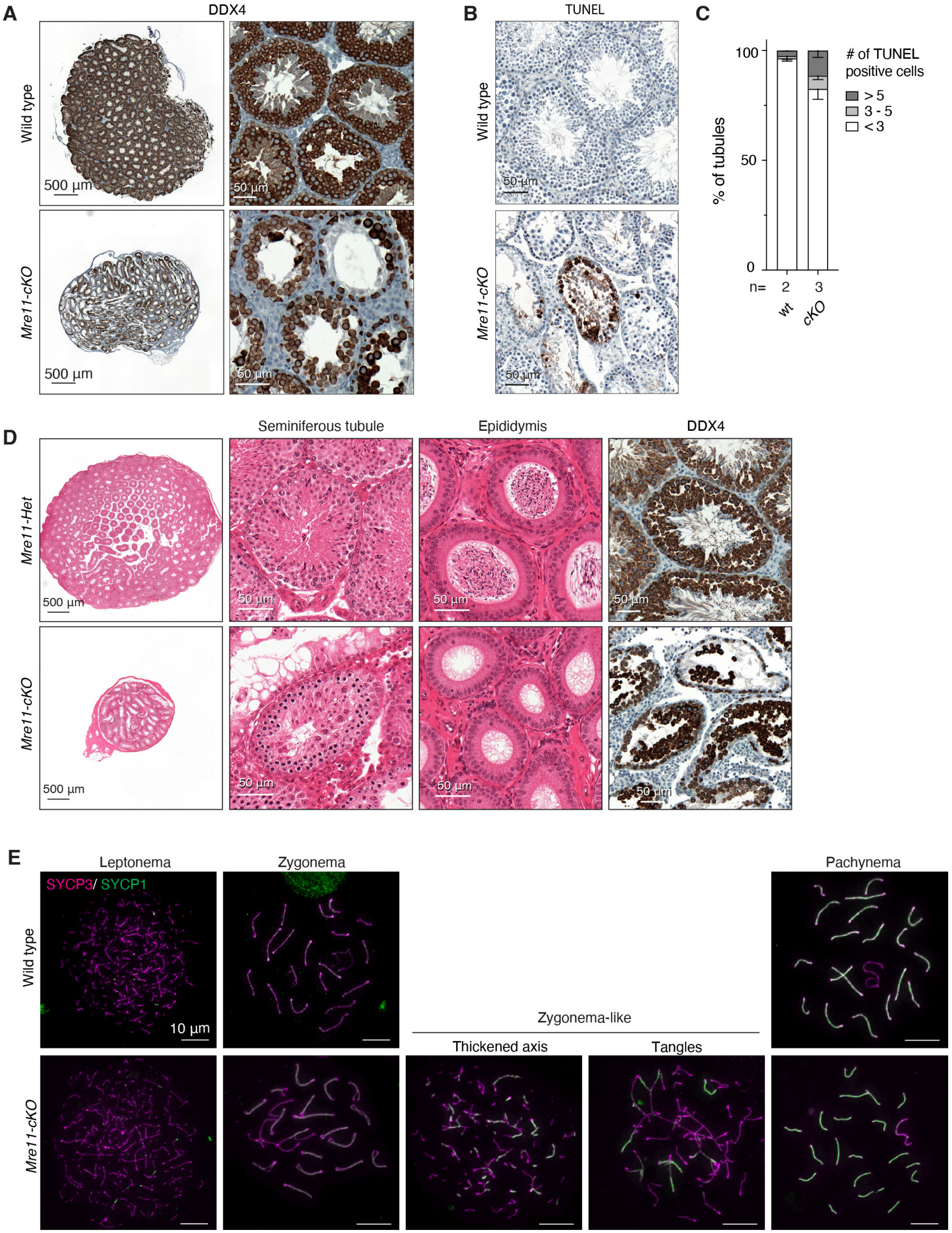


**Supplementary Figure 3. Spermatogenesis defects in *Mre11*-deficient mice**
(A) PFA-fixed testis sections at 7 wk of age stained for germ-cell marker DDX4.

(B) PFA-fixed seminiferous tubule sections at 7 wk of age stained with TUNEL to show apoptotic cells.
(C) Frequencies of tubules with the indicated number of TUNEL-positive cells. Error bars indicate mean ± range for the indicated number of animals.
(D) Bouin's fixed seminiferous tubule and epididymis sections stained with H&E (first three columns) and PFA-fixed seminiferous tubule sections stained for DDX4 (right column). Animals were 16 wk old.
(E) Representative spermatocyte spreads stained for SYCP3 and SYCP1.

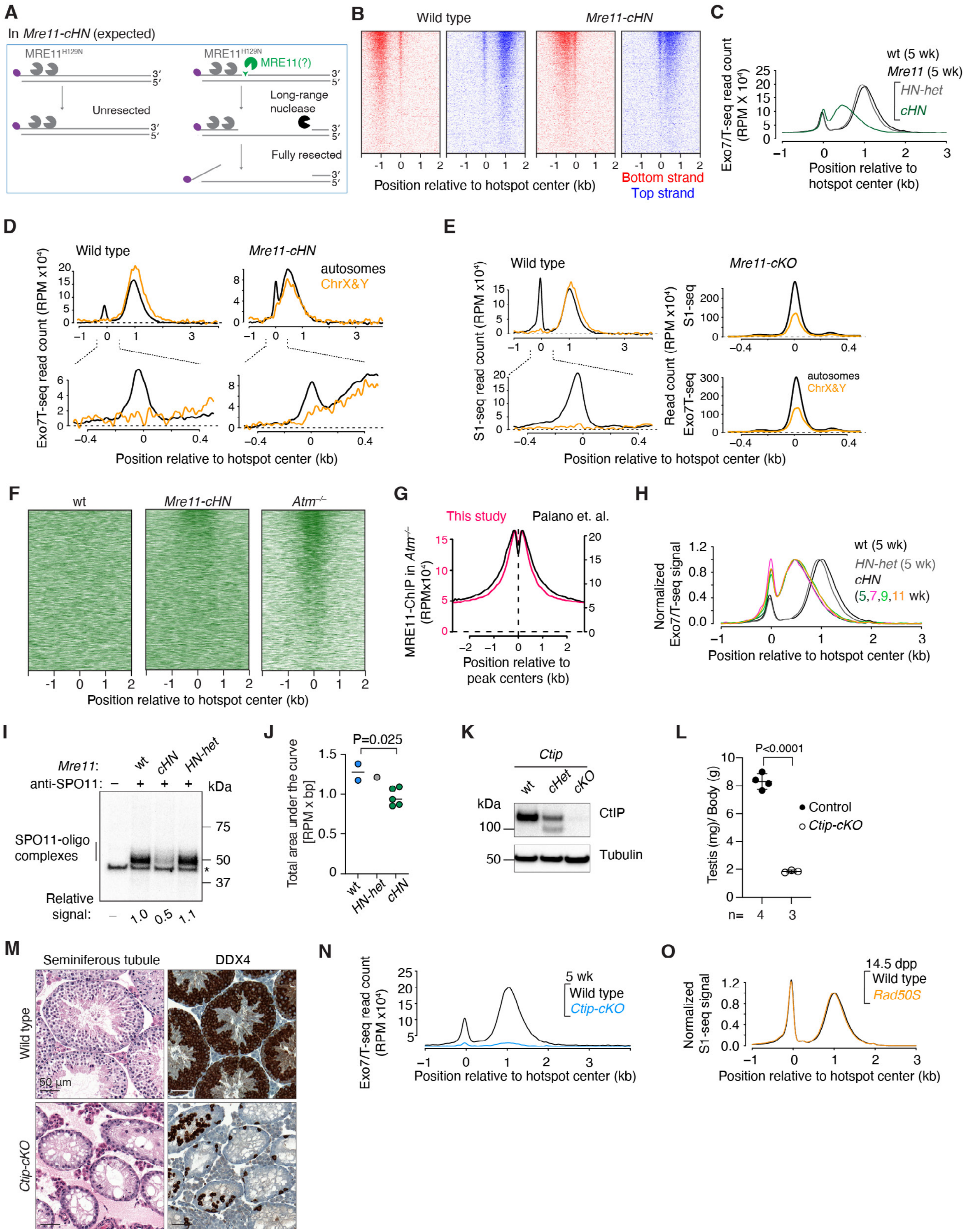


**Supplementary Figure 4. Phenotypes of *Mre11-cHN*, conditionally deleted *Ctip*, and *Rad50S*.**

(A) Schematic illustration of the originally anticipated DSB resection defects in *Mre11-cHN*. If the wild-type MRE11 protein is fully depleted by the time DSBs are formed and MRE11 is the only endonuclease that can initiate resection, then DSBs should remain unresected in *Mre11-cHN* (left). Alternatively, if residual wild-type MRE11 protein is retained for long enough after Cre-mediated excision to initiate resection, and if MRE11 is only involved in that initiation step, then any DSBs that are resected will be processed to the normal extent (right). Counter to these expectations, however, *Mre11-cHN* mice showed only a small apparent fraction of unresected DSBs plus a large population of resected DSBs with processing lengths that were much shorter than normal.
(B) Heatmaps of Exo7/T-seq signals around DSB hotpots from 5-wk-old mice. Two biological replicates each for wild type and *Mre11-cHN* were averaged.
(C) The non-normalized plot of **Figure 4D**.
(D) Exo7/T-seq signals from autosomal and sex chromosome hotspots. The profiles were generated by subtracting the Exo7/T-seq signal obtained in the 14.5-dpp *Spo11*$^{-/-}$ mutant and then averaging top and bottom strand reads after co-orienting them around hotspot centers. Data are smoothed with a 151-bp Hann window. The bottom panels show zoomed views (smoothed with a 51-bp Hann window) into the region around hotspot centers.
(E) Distinct patterns at hotspot centers for autosomal vs. sex chromosome hotspots depending on whether DSB resection is initiated. Left, S1-seq signals from autosomal and sex chromosome hotspots in wild type (14.5 dpp). In these mice, all DSBs are resected and the central signal (which is only from recombination intermediates) is seen only on autosomes. Right, S1-seq and Exo7/T-seq signals at hotspot centers in *Mre11-cKO* (5 wk old). In these mice, essentially all of the DSBs remain unresected and the central signal (which is now from unresected DSBs) is seen on both autosomes and sex chromosomes. Plots were generated as described in panel D, except that S1-seq signals from 14.5-dpp *Spo11*$^{-/-}$ mice were used for background subtraction of the S1-seq profiles.
(F) Heatmaps (data in 40-bp bins) of MRE11 ChIP-seq signals around DSB hotspots. Each line is a hotspot, ranked from strongest at the top (based on SPO11-oligo read count). The ChIP-seq signal at each hotspot was locally normalized by dividing by the total signal in a 4001-bp window around that hotspot's center. Each hotspot thus has a total value of 1, so that spatial patterns can be compared between hotspots of different strengths. Note that the central enrichment of MRE11 ChIP-seq signal is more pronounced the stronger the hotspot is.
(G) Good agreement between this study and previous data[14] for MRE11 ChIP-seq coverage around hotspots in *Atm*$^{-/-}$ mice.
(H) Reproducibility of resection profiles across a range of young adult ages for *Mre11-cHN* mice. Data are smoothed with a 151-bp Hann window and normalized to the peak height of resection endpoints. Resection profiles of wild type and 5 wk old *Mre11 HN-het* or *Mre11-cHN* are reproduced from **Figure 4D**. The profiles for 7, 9, and 11 wk old *HN-cHN* were each from a single library.
(I) Biological replicate of the experiment in **Figure 4F**, showing that both the reduction in the number of SPO11-oligo complexes and their altered electrophoretic mobility are reproducible in *Mre11-cHN* mice.
(J) Comparison of Exo7/T-seq signal strengths for wild type (n=2; 5 wk old) versus *Mre11 HN-het* (n=1; 5 wk old) or *Mre11-cHN* (n=5; 5–11 wk old) quantified as in **Figure 2B**. The P value is from a two-tailed Student's t test.
(K–N) Premeiotic depletion of spermatogonia in *Ctip* conditional deletion mice (*Ctip-cKO*). CtIP is an essential cofactor that promotes MRN nuclease activity[3] and its yeast orthologs (Sae2 in *S. cerevisiae* and Ctp1 in *S. pombe*) are needed for meiotic DSB resection[4,5,92,93,118]. Because *Ctip* is essential for viability, we attempted to evaluate its function in meiosis by combining a floxed *Ctip* allele[101,102] with *Ngn3-Cre*.

(K) Drastic depletion of CtIP protein in immunoblots of whole-testis extracts from *Ctip-cKO* mice (5 wk old). The faster migrating band in samples from *Ctip-cHet* (*Ctip*$^{wt/flox}$ *Ngn3-Cre*$^{+}$) or *Ctip-cKO* (*Ctip*$^{flox/–}$ *Ngn3-Cre*$^{+}$) may have originated from an internal translation start site on the transcript from the deleted allele; if so, the truncated form of CtIP appears to be stable only in the presence of the full-length protein.
(L) Reduced testis size in *Ctip-cKO* (5–8 wk old). Error bars indicate mean ± SD. The P value is from a Student's t test.
(M) Germ cell depletion in *Ctip-cKO*. Left, Bouin's fixed, H&E-stained seminiferous tubule sections (7 wk old). Right, PFA-fixed sections stained for DDX4 to mark germ cells. Note that most of the tubules had few or no DDX4-positive cells, unlike in *Mre11-cKO* (compare with **Figure S3A**). We conclude that CtIP is essential for germ cell maintenance in the mouse testis.
(N) Average Exo7/T-seq profiles at hotspots. Data are smoothed with a 151-bp Hann window. We observed only extremely weak Exo7/T-seq signals with resection tracts similar to wild type. We infer that this residual signal comes from a small fraction of phenotypically normal germ cells that escaped Cre-mediated excision. The non-normalized wild-type profile is reproduced from panel C.
(O) Normal resection in mice homozygous for *Rad50S* (K22M mutation; 14.5 dpp). Averaged S1-seq profiles around hotspots are shown. The *Rad50S* samples are indistinguishable from wild type, consistent with a previous report that this mutant has unperturbed meiotic progression[21]. Wild type is reproduced from **Figure 2A.** Data are smoothed with a 151-bp Hann window and normalized to the peak height of resection endpoints as described in **Figure 4D**.

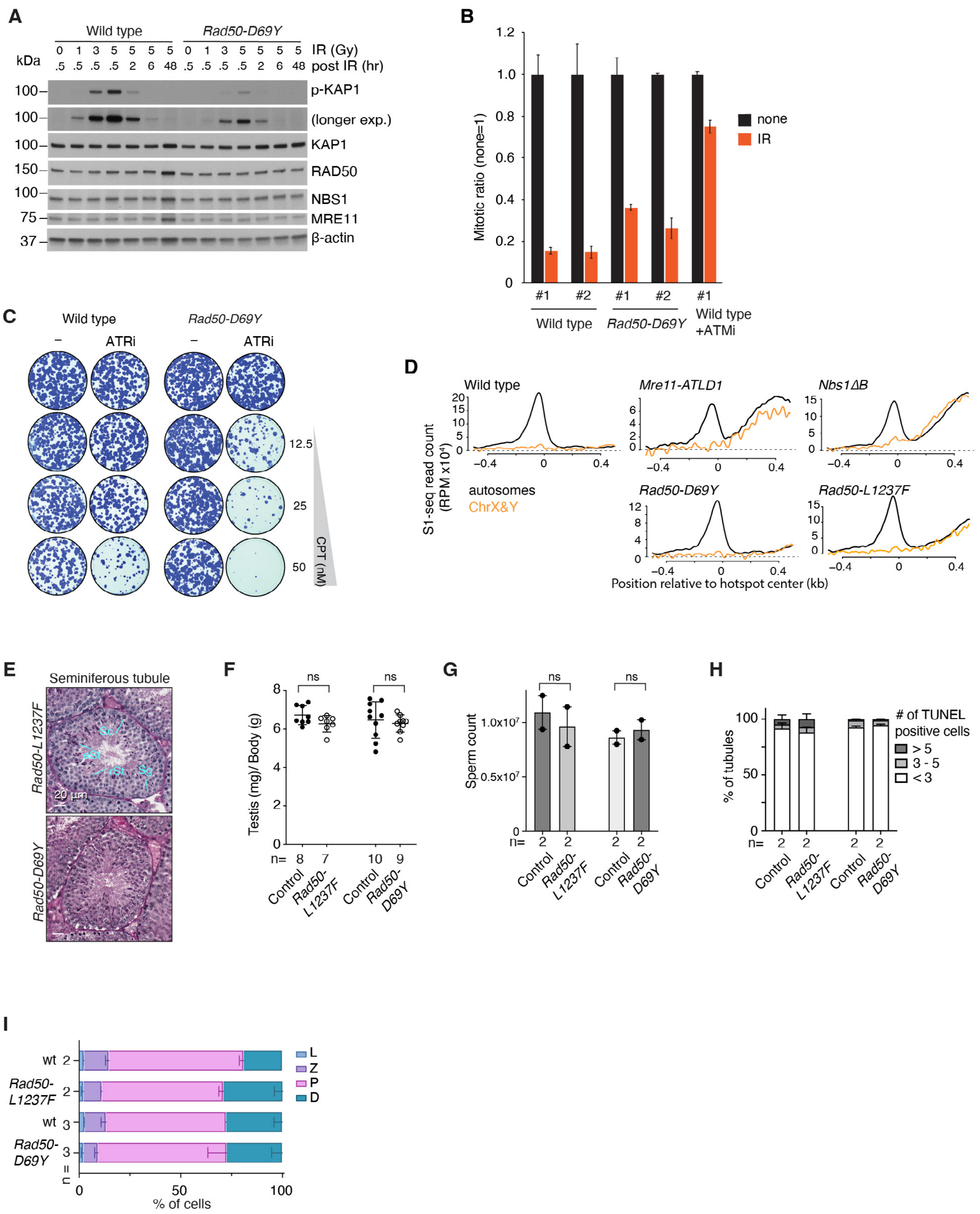


**Supplementary Figure 5. Mitotic and meiotic phenotypes of MRN hypomorphic mutants**
(A) Reduced ionizing radiation (IR)-provoked ATM signaling in *Rad50-D69Y* MEFs. MEFs from wild type and *Rad50-D69Y* mice were treated with different doses of IR, then cell-free extracts were prepared after various recovery times and ATM signaling was assessed by immunoblotting for phosphorylation of KAP1 on Ser-824. Phosphorylation was already evident at 30 min

following a 1-Gy IR exposure in wild-type cells, but was only detected at sharply reduced levels 30 min after an even higher dose (3 Gy) in the mutant. Note that levels of RAD50, MRE11, and NBS1 proteins were normal in the mutant, indicating that the mutation does not destabilize RAD50 or interfere with formation of the MRN complex.
(B) Reduced G2/M cell cycle checkpoint in *Rad50-D69Y* MEFs. After 1 hr recovery following irradiation with 3 Gy of IR, mitotic cells were quantified by measuring mitosis-specific phosphorylation of histone H3 Ser10 by flow cytometry. Pretreatment with an ATM inhibitor (KU55933, 10 μM) before irradiation served as an *Atm*-deficient control[119]. The mitotic indices of two independent *Rad50-D69Y* MEF lines were twofold higher than those of wild-type cells, consistent with reduced ATM activity. Error bars indicate means ± SD of three replicates.
(C) Increased DNA damage sensitivity in *Rad50-D69Y* MEFs when ATR is inhibited. Wild-type and *Rad50-D69Y* MEFs were treated with increasing doses of camptothecin (CPT) with and without concomitant inhibition of ATR (ATRi) with 50 nM VE822. In the absence of ATRi, the mutant showed no increase in CPT sensitivity compared to wild type. Wild-type cells were modestly more sensitive to CPT when ATR was inhibited than without ATRi, but the *Rad50-D69Y* mutant cells were substantially more CPT-hypersensitive in the presence of ATRi. These findings are consistent with the mutant cells being less able to compensate for ATR loss because they fail to activate ATM normally.
(D) S1-seq signals from autosomal and sex chromosome hotspots, from 14.5dpp mice, plotted as described in **Figure S4E**. Wild type is reproduced from **Figure S4E.**
(E) Grossly normal spermatogenesis in *Rad50-L1237F* or *Rad50-D69Y* mice. Bouin's fixed testis sections (9-11 wk of age) were stained with PAS (Periodic Acid-Schiff)-Harris Hematoxylin.
(F,G) Normal testis sizes (F) and sperm counts (G) in *Rad50-L1237F* or *Rad50-D69Y* mice. Error bars indicate mean ± SD. The results of Student's t tests are shown.
(H,I) Quantification of apoptotic tubules (H) and spermatocyte stages (I) based on SYCP3 staining. Error bars indicate mean ± range.

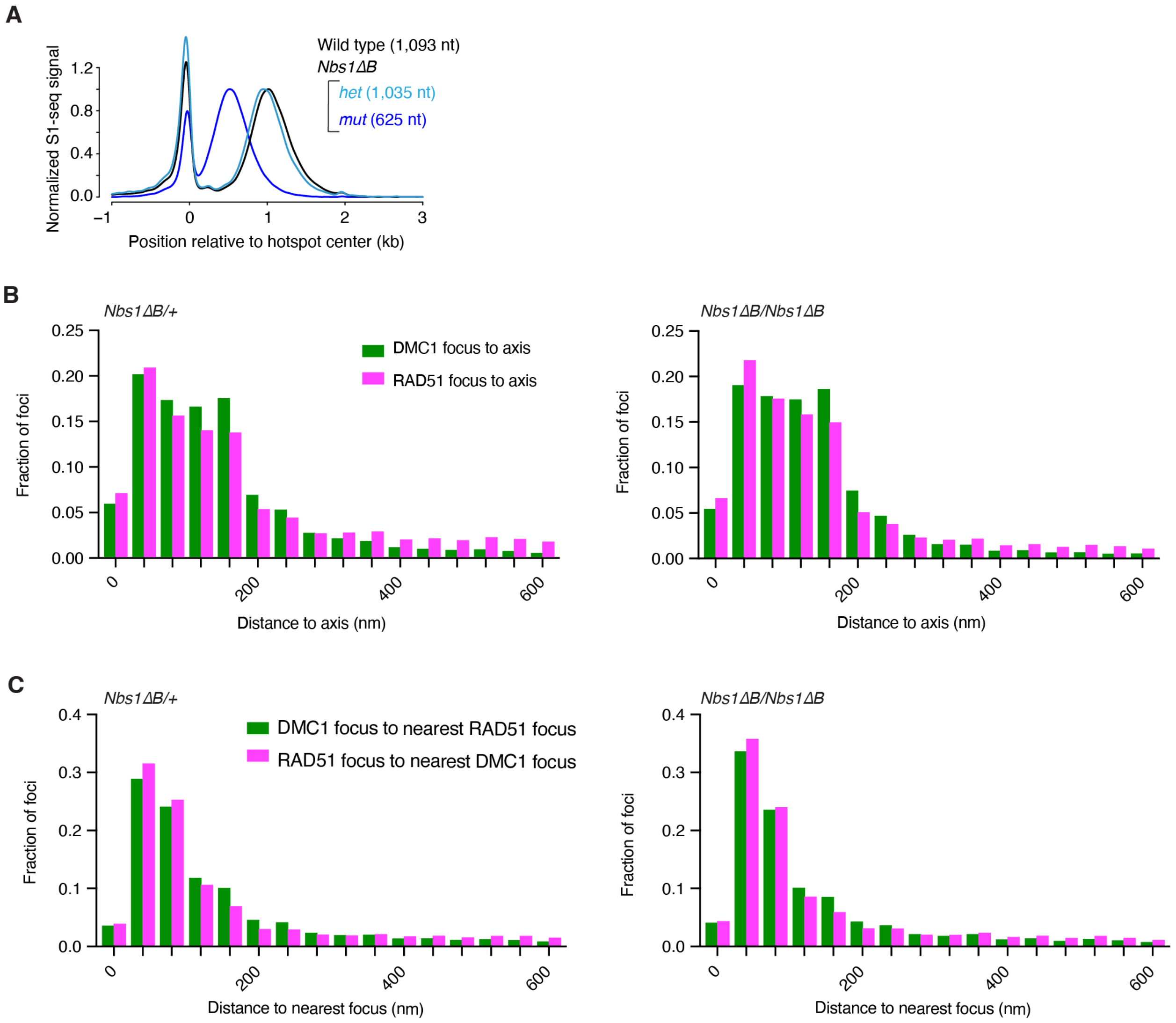


**Supplementary Figure 6. Spatial disposition of DMC1 and RAD51 foci in *Nbs1ΔB* spermatocytes**

(A) Genome-average S1-seq patterns at hotspots. Data from 14.5-dpp animals are smoothed with a 151-bp Hann window. Wild type and *Nbs1ΔB* homozygote (*mut*) profiles are reproduced from **Figure 5D**. Two replicates were averaged for *Nbs1ΔB* heterozygote or homozygote (*mut*) and three for wild type. Mean resection lengths are indicated in parentheses.

(B) The distance to the nearest axis (data in 40-nm bins) of DMC1 or RAD51 foci measured on SIM micrographs of spread spermatocytes. Distances larger than 600 nm were excluded for plotting purposes.

(C) Interfocus distances (data in 40-nm bins). Distances larger than 600 nm were excluded for plotting purposes.

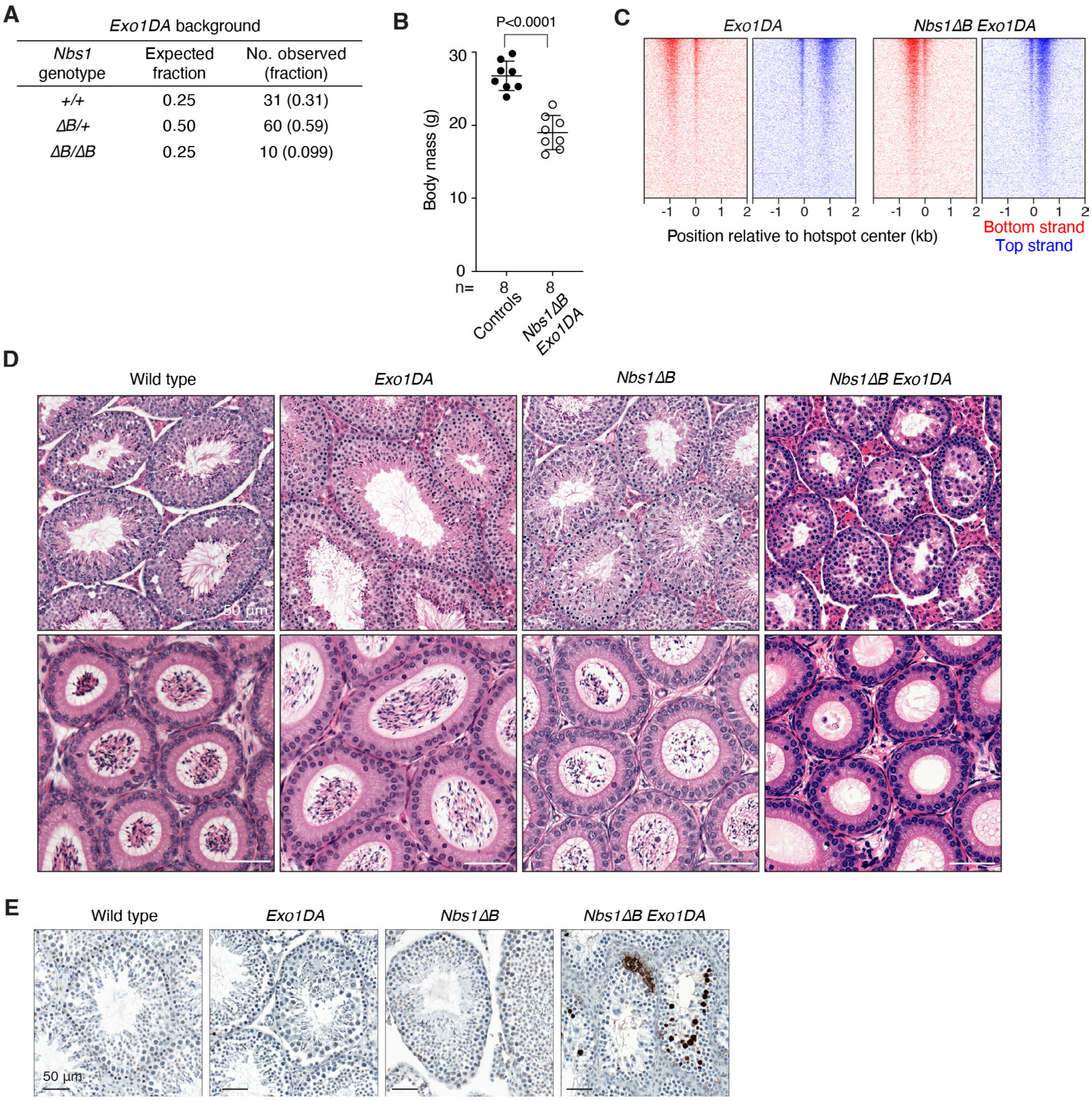


**Supplementary Figure 7. Phenotypes of *Nbs1ΔB Exo1DA* double mutants.**
(A) Submendelian recovery of *Nbs1ΔB Exo1DA* double mutant weanlings from mating of *Nbs1*$^{\Delta B/+}$ *Exo1*$^{DA/DA}$ animals. The recovery of double-mutant animals was significantly lower than expected (P = 0.0021, chi square goodness-of-fit test).
(B) Reduced body size in double mutant *Nbs1ΔB Exo1DA* animals compared with littermate controls. Error bars indicate mean ± SD. The P value is from a Student's t test.
(C) Stereotyped distribution of S1-seq signals around DSB hotspots from 16.5-dpp *Exo1DA* single mutants or *Nbs1ΔB Exo1DA* double mutants.
(D) Defective spermatogenesis in *Nbs1ΔB Exo1DA* double mutants. Bouin's fixed seminiferous tubule and epididymis sections of 7-wk-old wild type or mutant animals were stained with H&E.
(E) PFA fixed seminiferous tubule sections stained with TUNEL to show apoptotic cells.

**Supplemental Table S1. Summary of SIM imaging**

| | ***Nbs1*** $^{\Delta B/+}$ | ***Nbs1*** $^{\Delta B/\Delta B}$ |
|---|---|---|
| Mice analyzed | 2 | 2 |
| Images captured | 115 | 100 |
| Images per prophase I stage | Lep/Zyg, 101<br>Pachytene, 14 | Lep/Zyg, 87<br>Pachytene, 13 |
| RAD51 foci analyzed | 24,647 | 31,331 |
| DMC1 foci analyzed | 22,692 | 30,477 |
| *Axis-associated RAD51 | 19,560 (79.4%) | 27,047 (86.3%) |
| Axis-associated DMC1 | 21,274 (93.8%) | 28,866 (94.7%) |
| **Co-foci (RAD51) | 77.4% | 82.3% |
| Co-foci (DMC1) | 80.7% | 84.8% |

*A RAD51 or DMC1 focus was deemed axis-associated if was within 450 nm of the nearest axis.

**Co-foci were defined as axis-associated foci that were within 320 nm of a focus of the other protein.

**Supplemental Table S2. S1-seq, Exo7/T-seq and MRE11-ChIP mapping statistics**

| Method, genotype and age | No. of reads | No. mapped | No. mapped uniquely | Ref |
|---|---|---|---|---|
| S1-seq, wild type, 14.5 dpp | 16,731,828 | 15,406,997 | 8,196,974 | * |
| S1-seq, wild type, 14.5 dpp | 50,305,987 | 45,746,907 | 17,453,617 | * |
| S1-seq, wild type, 14.5 dpp | 23,836,476 | 21,804,210 | 3,206,542 | * |
| S1-seq, wild type, 14.5 dpp | 13,538,465 | 11,245,451 | 7,052,876 | * |
| S1-seq, wild type, 14.5 dpp | 20,563,792 | 18,317,528 | 8,721,466 | * |
| Exo7/T-seq, wild type, 14.5 dpp | 19,157,375 | 17,410,195 | 2,407,750 | * |
| Exo7/T-seq, wild type, 14.5 dpp | 9,625,692 | 8,364,663 | 1,786,410 | * |
| S1-seq, wild type, 11 wk | 11,950,314 | 9,289,056 | 5,281,339 | * |
| Exo7/T-seq, wild type, 5 wk | 23,860,774 | 20,240,541 | 4,400,942 | * |
| Exo7/T-seq, wild type, 5 wk | 22,254,127 | 20,189,150 | 4,348,064 | * |
| S1-seq, *Spo11*$^{-/-}$, 14.5 dpp | 22,406,609 | 19,888,057 | 7,898,087 | * |
| S1-seq, *Spo11*$^{-/-}$, 14.5 dpp | 9,557,181 | 7,472,470 | 3,751,662 | * |
| Exo7/T-seq, *Spo11*$^{-/-}$, 14.5 dpp | 18,057,775 | 16,654,402 | 9,993,210 | ** |
| Exo7/T-seq, *Spo11*$^{-/-}$, 14.5 dpp | 15,520,042 | 14,331,587 | 8,247,128 | ** |
| S1-seq, *Mre11-cKO*, 14.5 dpp | 9,253,463 | 7,394,058 | 3,682,713 | † |
| S1-seq, *Mre11-cKO*, 5 wk | 14,307,348 | 11,992,935 | 6,414,121 | † |
| S1-seq, *Mre11-cKO*, 5 wk | 12,932,363 | 11,316,956 | 5,849,802 | † |
| Exo7/T-seq, *Mre11-cKO*, 5 wk | 8,624,714 | 7,430,200 | 3,501,289 | † |
| Exo7/T-seq, *Mre11-cKO*, 5 wk | 11,837,141 | 9,949,034 | 3,500,795 | † |
| S1-seq, *Mre11-cHN*, 14.5 dpp | 22,798,321 | 20,066,591 | 6,546,369 | † |
| Exo7/T-seq, *Mre11-cHN*, 5 wk | 21,053,681 | 19,067,766 | 5,430,126 | † |
| Exo7/T-seq, *Mre11-cHN*, 5 wk | 14,536,250 | 12,405,858 | 5,020,437 | † |
| Exo7/T-seq, *Mre11-cHN*, 7 wk | 8,624,714 | 7,430,200 | 3,501,289 | † |
| Exo7/T-seq, *Mre11-cHN*, 9 wk | 19,594,185 | 17,972,385 | 5,128,521 | † |
| Exo7/T-seq, *Mre11-cHN*, 11 wk | 18,155,410 | 16,757,982 | 5,867,736 | † |
| Exo7/T-seq, *Mre11-HN het*, 5 wk | 19,865,984 | 18,103,199 | 5,278,163 | † |
| S1-seq, *Mre11-ATLD1*, 14.5 dpp | 13,140,150 | 11,101,025 | 2,661,923 | † |

| | | | | |
|---|---|---|---|---|
| **S1-seq, *Mre11-ATLD1*, 14.5 dpp** | 12,282,768 | 11,189,730 | 6,240,642 | † |
| **S1-seq, *Rad50-L1237F*, 14.5 dpp** | 33,428,256 | 30,063,707 | 19,641,976 | † |
| **S1-seq, *Rad50-L1237F*, 14.5 dpp** | 10,388,368 | 8,266,451 | 3,417,215 | † |
| **S1-seq, *Rad50-D69Y*, 14.5 dpp** | 38,599,640 | 34,017,973 | 17,269,962 | † |
| **S1-seq, *Rad50-D69Y*, 14.5 dpp** | 9,774,402 | 7,713,491 | 4,346,239 | † |
| **S1-seq, *Nbs1ΔB*, 14.5 dpp** | 32,029,388 | 28,791,187 | 16,807,532 | † |
| **S1-seq, *Nbs1ΔB*, 14.5 dpp** | 18,887,855 | 16,916,545 | 8,483,822 | † |
| **S1-seq, *Nbs1$^{ΔB/+}$*, 14.5 dpp** | 11,668,010 | 9,643,329 | 2,498,395 | † |
| **S1-seq, *Nbs1$^{ΔB/+}$*, 14.5 dpp** | 9,902,327 | 7,677,423 | 3,750,832 | † |
| **S1-seq, *Nbs1ΔB*, 16.5 dpp** | 7,999,728 | 6,272,866 | 3,014,706 | † |
| **S1-seq, *Nbs1ΔB*, 16.5 dpp** | 11,373,014 | 9,728,431 | 5,412,054 | † |
| **S1-seq, *Nbs1ΔB Exo1DA*, 16.5 dpp** | 12,512,577 | 11,460,477 | 6,225,902 | † |
| **S1-seq, *Nbs1ΔB Exo1DA*, 16.5 dpp** | 9,300,830 | 7,216,734 | 3,609,598 | † |
| **S1-seq, *Exo1DA*, 16.5 dpp** | 12,227,344 | 11,177,166 | 6,319,057 | † |
| **S1-seq, *Exo1DA*, 16.5 dpp** | 27,628,884 | 24,038,637 | 6,243,144 | † |
| **S1-seq, *Rad50S*, 16.5 dpp** | 22,733,841 | 20,105,504 | 12,218,008 | † |
| **S1-seq, *Rad50S*, 16.5 dpp** | 12,066,801 | 10,861,659 | 6,378,596 | † |
| **S1-seq, *Ctip-cKO*, 14.5 dpp** | 24,539,681 | 21,274,883 | 4,776,736 | † |
| **Exo7/T-seq, *Ctip-cKO*, 5 wk** | 19,846,982 | 17,326,860 | 6,027,926 | † |
| **S1-seq, wild type, 12.5 dpp** | 55,422,810 | 50,863,055 | 15,658,589 | * |
| **S1-seq, wild type, 13.5 dpp** | 58,845,411 | 54,411,337 | 17,560,563 | * |
| **S1-seq, wild type, 15.5 dpp** | 51,945,205 | 48,042,167 | 24,049,978 | * |
| **S1-seq, wild type, 16.5 dpp** | 54,938,845 | 49,928,756 | 14,745,494 | * |
| **S1-seq, wild type, 12.5 dpp** | 22,751,133 | 20,356,990 | 3,700,654 | † |
| **S1-seq, wild type, 13.5 dpp** | 24,898,561 | 22,742,274 | 3,215,326 | † |
| **S1-seq, wild type, 15.5 dpp** | 24,872,788 | 22,686,216 | 3,842,106 | * |
| **S1-seq, wild type, 16.5 dpp** | 25,804,707 | 23,468,594 | 4,398,765 | * |
| **S1-seq, wild type, 12.5 dpp** | 37,003,923 | 7,455,456 | 1,965,592 | † |
| **S1-seq, wild type, 13.5 dpp** | 31,228,109 | 6,175,560 | 1,658,463 | † |
| **S1-seq, wild type, 14.5 dpp** | 25,610,550 | 5,155,404 | 1,151,530 | † |
| **S1-seq, wild type, 15.5 dpp** | 31,938,187 | 6,541,586 | 1,372,672 | † |

| | | | | |
|---|---|---|---|---|
| **S1-seq, wild type, 16.5 dpp** | 32,078,661 | 6,365,300 | 1,388,773 | † |
| **MRE11-ChIP, wild type, 7wk** | 11,853,063 | 10,483,250 | 4,566,646 | † |
| **MRE11-ChIP, wild type, 7wk** | 14,794,788 | 14,359,551 | 7,658,364 | † |
| **MRE11-ChIP, *Mre11-cHN*, 7wk** | 9,273,480 | 8,823,644 | 4,457,326 | † |
| **MRE11-ChIP, *Mre11-cHN*, 7wk** | 12,186,242 | 11,649,413 | 5,486,836 | † |
| **MRE11-ChIP, *Mre11-cHN*, 7wk** | 10,574,081 | 10,015,905 | 4,803,917 | † |
| **MRE11-ChIP, *Atm*$^{-/-}$, 7wk** | 11,801,432 | 10,602,025 | 4,011,579 | † |

* Kim et al. 2024[32]
** Xu et al. 2023[114]
† This study

**Supplemental Table S3. Antibodies.**

| Primary Antibodies | | | | | |
|---|---|---|---|---|---|
| Target | Dilution for IF | Dilution for IB | Host Species | Supplier | Catalog number or reference |
| SYCP3 | 1:400 or 1:200 | n/a | Mouse | Santa Cruz | sc-74569 |
| SYCP1 | 1:200 | n/a | Rabbit | Abcam | ab15090 |
| RPA2 | 1:1000 | n/a | Rabbit | Abcam | ab76420 |
| DMC1 | 1:100 | n/a | Rabbit | Santa Cruz | sc-22768 |
| DMC1 | 1:50 | n/a | Guinea pig | home-made T113-1, ABclonal Biotechnology | Ref.[64] |
| RAD51 | 1:50 | n/a | Rabbit | Santa Cruz | sc-8349 |
| γH2AX | 1:6000 | n/a | Rabbit | Abcam | ab2893 |
| α-tubulin | n/a | 1:5000 | Rat | Bio-Rad | MCA78G |
| MRE11 | n/a | 1:2000 | Rabbit | Novus Biologicals | NB100-142 |
| MRE11 | n/a | 1:5000 | Rabbit | Custom made (Petrini laboratory) | Ref.[22] |
| RAD50 | n/a | 1:2000 | Rabbit | Novus Biologicals | NB100-154 |
| RAD50 | n/a | 1:5000 | Rabbit | Custom made (Petrini laboratory) | Ref.[22] |
| NBS1 | n/a | 1:2000 | Rabbit | Novus Biologicals | NB100-143 |
| NBS1 | n/a | 1:5000 | Rabbit | Custom made (Petrini laboratory) | Ref.[22] |
| KAP1 | n/a | 1:5000 | Mouse | Santa Cruz | sc-136102 |
| pKAP1 (Ser824) | n/a | 1:5000 | Rabbit | Abcam | ab70369 |
| β-ACTIN | n/a | 1:40000 | Mouse | Abcam | ab49900 |
| | Dilution for Flow cytometry | | | | |
| pHistone H3 (Ser10) | 1:200 | | Rabbit | Millipore | 06-570 |
| | Dilution for IP | | | | |
| SPO11-180 | 1:2000 | | mouse | MSKCC Antibody and Bioresource Core Facility | n/a |
| | Dilution for Histology | | | | |
| MRE11 | 1:5000 | | Rabbit | Novus Biologicals | NB100-142 |
| DDX4 | 0.17 µg/ml | | Rabbit | Abcam | ab13840 |

| | Dilution for ChIP | | | |
|---|---|---|---|---|
| MRE11 | 10 µl/ reaction | Rabbit | Novus Biologicals | NB100-142 |
| DMC1 | 20 µl/ reaction | Mouse | Abcam | ab11054 |
| RAD51 | 10 µl/ reaction | Mouse | Novus Biologicals | NB100-148 |
| RPA2 | 10 µl/ reaction | Rabbit | Abcam | ab76420 |

**Secondary Antibodies**

| Target | Dilution for IF | Fluorophore | Host Species | Supplier | Catalog number |
|---|---|---|---|---|---|
| anti-Guinea pig | 1:100 | Alexa-488 | Goat | Invitrogen | A-11073 |
| anti-Mouse | 1:200 | CF405S | Goat | Biotium | BT20080 |
| anti-Mouse | 1:400 | Alexa-594 | Goat | Invitrogen | A-11005 |
| anti-Rabbit | 1:100 or 1:400 | Alexa-555 | Donkey | Invitrogen | A-31572 |
